# mRNArchitect: sequence design of mRNA medicines

**DOI:** 10.1101/2024.12.03.626696

**Authors:** Helen M. Gunter, Alexandra Balicki, Magdalena Budzinska, Jonathon Waterhouse, Theodore Leonard, Yi Peng, Abhishek Kulkarni, Mohammad Zardbani, Michelle Sidharta, Jessica Morrison, Bettina Dietmair, Aayushi Ghodasara, Sneha Lakshman Das, Seth W. Cheetham, Natasha Chaudhary, Tim R. Mercer

## Abstract

Sequence design is a critical first step in the development of mRNA medicines. The primary sequence can be optimised to improve translation, stability and immunogenicity. Here, we present *mRNArchitect*, a software toolkit for the design and optimisation of primary mRNA sequences according to codon usage, secondary structure and manufacturability. *mRNArchitect* supports multiple optimisation strategies tailored for short or long expression profiles, and different clinical applications, including vaccines, cell and gene therapies. *mRNArchitect* is complemented by a catalogue of validated RNA elements, including UTRs, microRNA binding sites, and regulatory elements, that provide programmable, cell-type-specific control of mRNA expression. We experimentally validate *mRNArchitect* designs, demonstrating improvements in expression, stability, and immunogenicity, across multiple cell types and tissues compared to current design methods. *mRNArchitect* is provided as an experimentally validated, software package for the design and optimisation of mRNA medicines, available at: https://basefacility.org.au/software/.

## INTRODUCTION

Sequence design is a key first step when developing a new mRNA medicine. The primary sequence determines the expression and stability of an mRNA throughout its life cycle. Optimising the sequence design can enhance the performance of an mRNA medicine by improving translation, increasing stability and reducing reactogenicity^1,2^.

mRNA medicines are being developed for an increasing range of clinical applications, including vaccines, protein replacement, gene editing, and cell therapies^3^. However, different clinical applications impose different challenges and opportunities for mRNA design. For example, mRNA vaccines may benefit from extensive secondary structure, which may incite the innate immune system and confer an adjuvant effect, whilst therapies that require frequent dosing should minimise these inflammatory properties^4^.

The design of an mRNA often aims to maximise the total output of target proteins. This total protein output is determined by both the translational strength and the stability of the mRNA sequence. The sequence can be enhanced for translation by selecting codons frequently used in recipient cells^5,6^, while the stability of the sequence can be increased with secondary structures^7,8^ (**Figure 1a**). Furthermore, selecting different untranslated regions (UTRs) can regulate both translation and stability in different cell types and tissues^9^.

**Figure 1.**
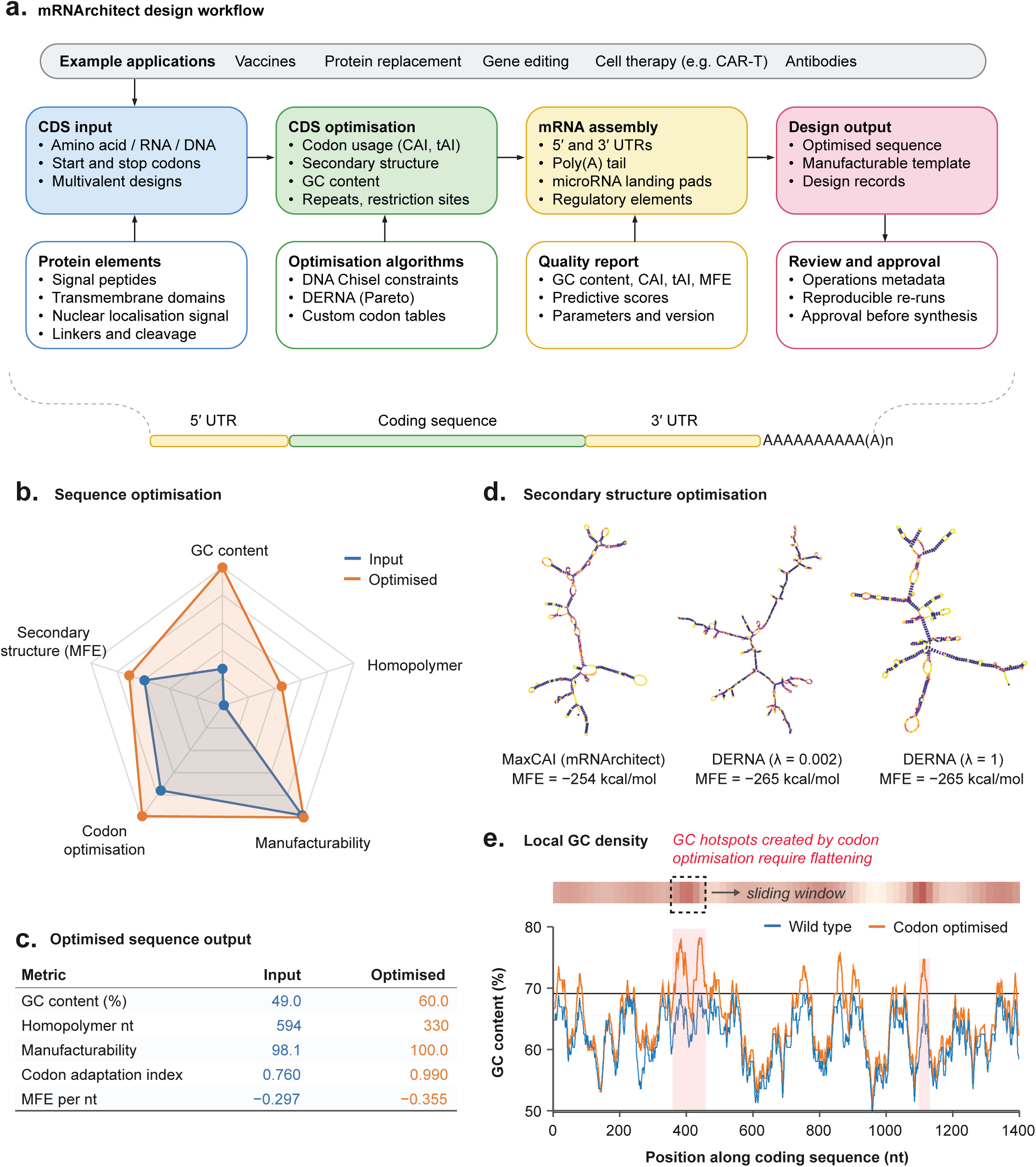
The *mRNArchitect* design workflow and its effect on sequence composition. (a) Design workflow. A coding sequence is supplied as amino acid, RNA or DNA, optionally with protein elements; the CDS is optimised for codon usage, secondary structure, GC content and sequence repeats using DNA Chisel constraints, DERNA or custom codon tables; UTRs, poly(A) tail and regulatory elements are then assembled, and a quality report and design record accompany the output. (b) Radar plot comparing an input sequence with its *mRNArchitect*-optimised counterpart across five design metrics, each scaled from 0 to 1. (c) The same metrics as absolute values. (d) Predicted secondary structures of one coding sequence optimised by MaxCAI and by DERNA at two λ settings, with minimum free energy below each. (e) Local GC content in a sliding window along the coding sequence, wild type against codon optimised. The heat strip above marks the GC-rich regions that optimisation flattens.

The expression of mRNA within the cell also presents engineering opportunities. Sequence elements can be incorporated into the mRNA to provide cell-type specific expression or modulate the subcellular localisation^10^. For example, microRNAs can regulate mRNA stability^11^, and signal peptides and transmembrane domains can direct endogenous processing, presentation, or secretion pathways^12^. mRNA medicines can encode large, multi-domain, or repetitive sequences. For example, chimeric antigen receptors, fusion proteins, and polyvalent vaccine antigens can encompass multiple functional domains, each of which may be translated, cleaved and processed within the cell^13^.

In addition to improving performance, mRNA medicines must be manufactured with good yield and quality. Effective manufacture requires a reliably synthesized DNA template that can be reliably *in vitro* transcribed into high-quality mRNA. GC-rich, structured or repetitive sequences can be difficult to synthesise, resulting in a poor-quality mRNA product, including high levels of truncated or damaged mRNA species that undermines mRNA performance. Accordingly, sequence design must also ensure the manufacturability of mRNA medicines.

To address these design considerations, we developed *mRNArchitect*, a software toolkit for the design of mRNA sequences. *mRNArchitect* supports different optimisation strategies. including codon optimisation and secondary structure and can be tailored to different clinical applications. *mRNArchitect* is also complemented by a catalogue of validated RNA elements that provide programmable, cell-type specific expression. We benchmark the *mRNArchitect* designs against alternative tools and methods and demonstrate the *in vitro* performance of our designs across multiple cell types and therapeutic applications.

## RESULTS

### Optimisation algorithm

We have developed *mRNArchitect*, software to optimise mRNA medicine sequences. The user supplies a coding sequence (CDS) encoding the protein of interest, in amino acid, RNA, or DNA format, which should initiate with a methionine codon (M, AUG) and terminate with one or more stop codon(s). The coding sequence may also include additional functional elements, such as signal peptides, nuclear localisation signals, or affinity tags, that are required for the protein’s correct processing or function. We provide a table of common, known and validated sequence protein domains in Supplementary Table 1.

To optimise the sequence, the users can select the preferred optimisation algorithm, whether to prioritise codon optimisation, which is recommended for most applications, or to prioritise secondary structure, which can increase the stability of an mRNA.

For codon optimisation, *mRNArchitect* uses *DNA Chisel*^14^ to iteratively select the preferred codons of the target organism according to either the Codon Adaptation Index (CAI)^5^, which favours codons enriched in housekeeping genes, or the tRNA Adaptation Index (tAI)^15^, which favour codons enriched in the tRNA pool of an organism. *mRNArchitect* can balance codon optimisation against other parameters (**Figure 1b**). Optimisation parameters in mRNArchitect are translated into DNA Chisel specifications, which are classified as either hard constraints or soft constraints (optimisation objectives). Hard constraints must be strictly satisfied and include GC content bounds and restriction site avoidance. In contrast, soft constraints define optimisation objectives whose scores are maximised, such as CAI/tAI and the avoidance of repeated k-mers (**Figure 1c**).

*mRNArchitect* can also be used to optimise secondary structure using DERNA^16^, which maximises intramolecular base-pairing in mRNA designs, resulting in a highly structured sequence (**Figure 1d**). Users may also balance secondary structure and codon optimisation using pareto-optimisation.

The manufacturability of the DNA template and mRNA product is another critical consideration during design, as sequences that are difficult to manufacture often produce low-quality mRNA products with compromised performance. GC content, which is correlated with stable secondary structures and human-optimised codons, must be managed to enable DNA manufacture and mRNA synthesis. In addition to imposing global constraints on GC content, *mRNArchitect* allows users to modulate GC content using a sliding window and resolve localised GC-rich regions that can be difficult to manufacture (**Figure 1e**). Hard constraints can also be used to exclude sequences, such as homopolymer tracts, stable hairpins, and restriction enzyme binding sites used in manufacture.

Many mRNA medicines, such as cell therapies and polyepitope vaccines, have a modular architecture, comprising multiple functional domains joined by linkers. To assist the optimisation of these modular mRNAs, *mRNArchitect* can independently optimise sub-regions of the provided sequence. This allows users to impose different optimisation parameters to different regions or leave specific regions unoptimized. For example, this can preserve validated linker sequences or reduce local secondary structure around the translation start site^17^.

### Running mRNArchitect

Following the optimisation of the input sequence, *mRNArchitect* generates a design report that includes descriptive statistics, including nucleotide composition, CAI and tAI scores, MFE and predicted secondary structure in dot-bracket notation. The design report can undergo review and approval prior to synthesis, with example design reports for a range of common mRNAs provided (**Table S2)**.

To benchmark the computational performance of *mRNArchitect*, we optimised 80 coding sequences from the human ORF dataset, with lengths ranging from 100–8000nt, using default and constrained codon optimisation parameters under a local installation (**see Methods**). With default parameters, optimisation time increased linearly with length, taking ∼1 second for ∼1000 nt constructs (**Figure S1a**). However, even for sequences of equal length, optimisation time varied due sequence complexity, with difficult sequences requiring an average of 4-fold more time to explore a larger solution space before identifying a solution. Constrained optimisation conditions resulted in a ∼5 x increase in optimisation time, consistent with the more restricted solution space. The peak memory usage increased approximately linearly with sequence length and was largely independent of the optimisation parameters **Figure S1b**).

### Optimisation of human genes using *mRNArchitect*

To investigate the optimisation of mRNA sequences using *mRNArchitect*, we assembled six sets of gene sequences to serve as representative datasets to assess sequence optimisation, including antibodies, antigens, CAR-T constructs, cancer-associated genes, housekeeping genes, and human open reading frames.

To quantify the degree of sequence remodelling by *mRNArchitect*, we calculated the Hamming distance ratio between wild-type and optimised sequences, defined as the proportion of nucleotide positions that differ in each pairwise comparison (**Figure S2a**). Wild-type cancer, housekeeping and human ORF sequences showed a mean ratio of 0.15–0.20 (range 0–0.372) following optimisation, corresponding to 15–20% of nucleotides being changed.

We next evaluated the optimised properties of the different mRNA sequences. As expected, sequence properties associated with hard or soft constraints converged tightly across the sequence datasets, following optimisation. The Codon Adaptation Index (CAI) increased across all gene classes approaching a CAI value of 1.0, with a corresponding reduction in rare codons (**Figure S2b**). Global GC content also converged upon a narrow range following optimisation (∼55–65%), indicating improved construct manufacturability.

A range of predictive models have been established to predict the functional performance of mRNA sequences. We applied predictive models of mRNA translation efficiency (RiboNN) and stability (CSC, DegScore) to the optimised sequences to predict the impact of optimisation on sequence performance^8,18,19^. Predicted measures of translation and stability scores were broadly improved following optimisation across all mRNA classes; however, these predicted functional gains were more modest and variable (**Figure S2b**).

### Benchmarking *mRNArchitect* to alternative tools

We next benchmarked *mRNArchitect* designs against other mRNA design tools. *mRNArchitect* was first used to optimise firefly luciferase (FLuc) and eGFP reporter mRNAs for expression in human cells using four strategies: codon optimisation by CAI (MaxCAI) or tAI (MaxtAI), maximal secondary structure by minimising MFE (MinMFE), and a Pareto-optimal balance of secondary structure and CAI (ParetoMFE) (**Figure 2a, Table S3**).

**Figure 2.**
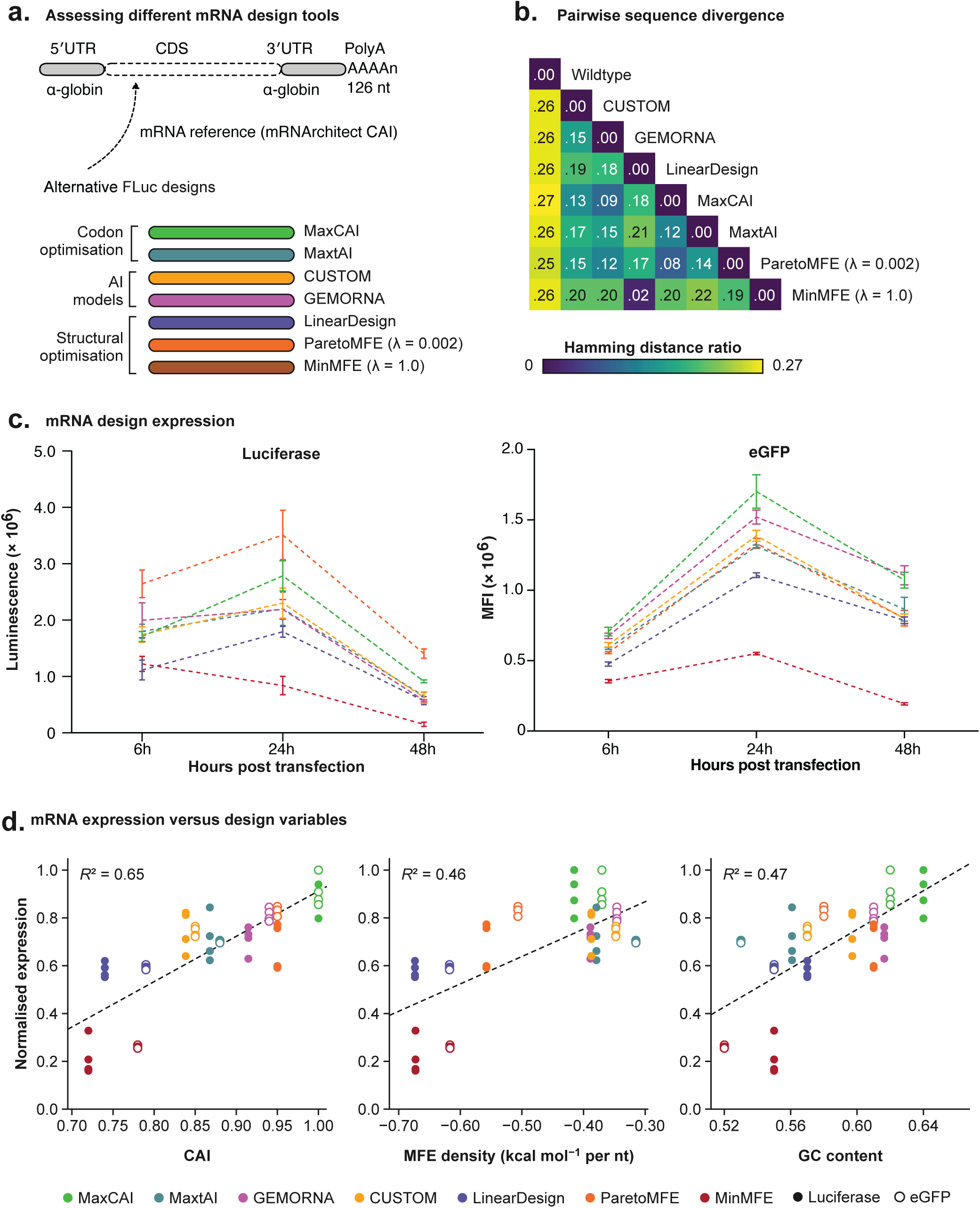
Benchmarking *mRNArchitect* against alternative mRNA design tools. (a) Constructs. A common human α-globin 5′ UTR, mouse α-globin 3′ UTR and 126 nt poly(A) tail were combined with eight alternative firefly luciferase coding sequences, grouped as codon optimisation (MaxCAI, MaxtAI), AI models (CUSTOM, GEMORNA) and structural optimisation (LinearDesign, ParetoMFE (λ = 0.002) and MinMFE (λ = 1.0)). (b) Pairwise Hamming distance ratio between the wild-type sequence and each design. (c) Luciferase luminescence and eGFP fluorescence at 6, 24 and 48 h post transfection. (d) Normalised reporter expression regressed against CAI, MFE density and GC content. Closed symbols, luciferase; open symbols, eGFP; dashed line, linear fit.

We next compared the *mRNArchitect* designs against sequences optimised by a range of alternative design tools. These include *LinearDesign*, which prioritises RNA secondary structure^2^, and three AI-based codon optimisation models: *GEMORNA*, *CodonTransformer* and *Codon Usage to Specific Tissue OptiMizer (CUSTOM)* ^20–22^.

To compare the degree of remodelling introduced by each tool, we calculated Hamming distance ratios (**Figure 2b**). Every tool remodelled the wild-type sequences (25–27% of nucleotides). However, divergence between the optimised designs was lower. GEMORNA and CUSTOM were closest to *mRNArchitect MaxCAI* (0.09 and 0.13, respectively) showing the convergence of AI models with codon optimisation. *LinearDesign* and *mRNArchitect* MinMFE (DERNA λ = 1.0) sequences were near-identical (0.02), forming similar secondary structures, while *mRNArchitect* ParetoMFE showed a secondary structure intermediate between MaxCAI and MinMFE (**Figure 4a, b**).

We also evaluated sequence features across all tools, including GC content, MFE, CAI, predicted half-life^8^, predicted translation (*RiboNN*)^19^, and Codon Stability Coefficient (*CSC*)^18^ (**Figure S3**). *LinearDesign* sequences showed the highest and most variable MFE and predicted half-life, consistent with prioritisation of stable secondary structures. GEMORNA sequences were the most variable of any tool. By contrast, *mRNArchitect’s* codon optimised designs (*MaxCAI, tAI*) produced the most consistent values across all six metrics.

### Expression of optimised mRNA sequences

We next compared the expression of *mRNArchitect’s* codon (CAI, tAI) and secondary structure based (MinMFE, ParetoMFE) optimisations to other design tools and strategies. We manufactured a subset of the optimised eGFP, and FLuc sequences generated by five different design tools (**Table S3**). Each mRNA was manufactured to high-quality control specifications, including sequence identity, integrity, purity, and yield (**Figure S4**). The mRNAs were then transfected into HEK293 cells and expression was measured at 6, 24 and 48 hours (**Figure 2c**).

For both reporter mRNAs, *mRNArchitect MaxCAI* produced the most consistently high expression. The effect was clearest at 24 hours, the time point of peak expression, and remained apparent at 48 hours. *mRNArchitect ParetoMFE* Luciferase also showed the highest expression of all Luciferase mRNAs, however *ParetoMFE* eGFP showed a more moderate expression, demonstrating sequence specific differences. These results support maximal CAI codon optimisation as an effective and reliable strategy for increasing mRNA expression. Among the remaining designs, *GEMORNA, CUSTOM* and *MaxtAI* performed similarly across time points and constructs, while *LinearDesign* and *MinMFE* showed lower expression.

Given that the different design tools generate mRNA sequences with diverse properties, we next examined the correlation between sequence properties and expression. CAI was positively correlated with normalised expression (R²=0.65) **Figure 2d**), while GC content and MFE were more moderately correlated with expression (R²=0.47 and R²=0.46 respectively). CAI explained the majority of the variance in expression for each fluorophore, in comparison to MFE density, GC content, tAI and DegScore predicted half-life (**Figure S5a, b**).

### In-vial stability of optimised mRNA sequences

In-vial stability is a key quality attribute in the manufacture of mRNA medicines, determining shelf-life and storage requirements, where degradation occurs primarily through hydrolysis and shearing^19^. Sequence design can mitigate hydrolysis by avoiding unstable codons^18^ and compacting structures that protect phosphodiester bonds from hydrolytic attack^2^. To benchmark the in-vial stability of *mRNArchitect*-designed mRNA against other design tools, we subjected *eGFP* and *FLuc* mRNAs to accelerated thermal degradation^23^ at 90°C and quantified their integrity by capillary gel electrophoresis at each timepoint (**Figure 3a**).

**Figure 3.**
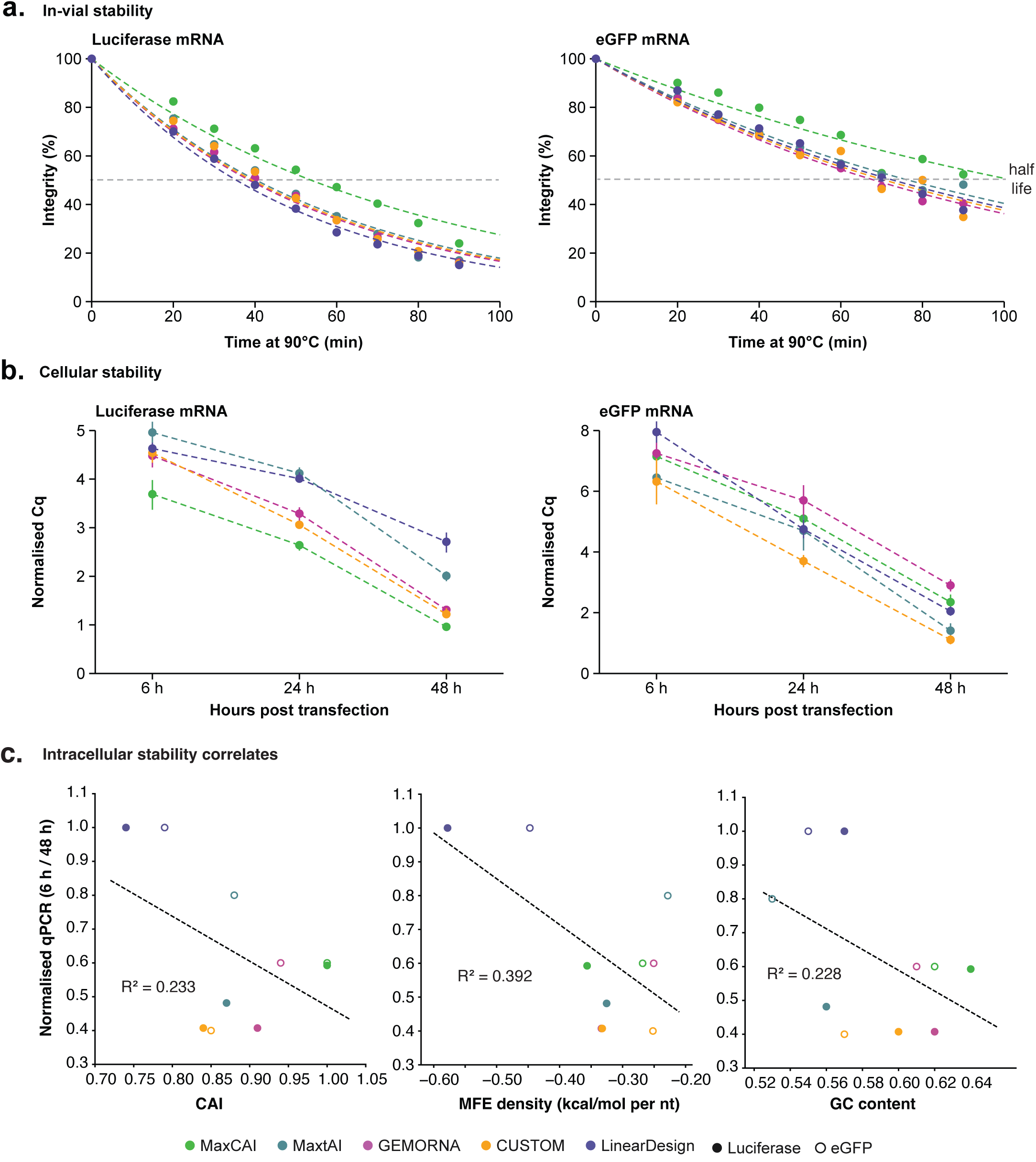
In-vial and intracellular stability of mRNA designs. (a) Remaining intact mRNA after accelerated thermal stress at 90 °C, measured by capillary gel electrophoresis, for luciferase and eGFP designs. Dashed lines are one-phase exponential fits. (b) Intracellular transcript abundance by RT-qPCR at 6, 24 and 48 h post transfection, normalised to GAPDH. (c) Intracellular mRNA persistence, the ratio of normalised mRNA abundance at 6 h to 48 h, regressed against CAI, MFE density and GC content. Closed symbols, luciferase; open symbols, eGFP.

For both reporters, *mRNArchitect MaxCAI* showed the greatest thermostability of the five design tools tested. The Luciferase designs retained the highest proportion of intact mRNA at 90 minutes (22.2% compared to 13.6–15.5% for the other tools), with a half-life of 54.6 minutes and a correspondingly lower decay constant (k=0.0127). A similar trend was observed for eGFP, demonstrating its generalisability. This greater thermostability likely reflects the higher GC content and low Uridine content of human-optimised codons, which confer a protective, compacting effect (**Figure S6a, b**).

### Intracellular stability of optimised mRNA sequences

Intracellular stability is a key determinant of mRNA medicine performance, since it governs the durability of expression after delivery, and is driven primarily by RNA processing pathways and nucleases. Therefore, we next tested the intracellular stability of mRNA designs generated using *mRNArchitect*, against designs from the other tools.

We transfected *eGFP* and *FLuc* mRNA designs into HEK293T cells and extracted RNA at 6, 24, and 48 hours post-transfection. The abundance of mRNA designs was then measured by RT-qPCR and normalised to *GAPDH*, with primer and probe efficiency and linearity were validated by standard curve prior to sample analysis (**see Methods**).

At 6 hours post-transfection, all five *eGFP* designs showed similar mRNA abundance relative to *GAPDH*, however, by 24 and 48 hours, mRNA abundance decreased sharply (**Figure 3b**). *LinearDesign* mRNAs showed the highest relative intracellular abundance at all timepoints, and the lowest change in mRNA abundance between 6 and 48h post transfection, indicating the greatest intracellular persistence. *mRNArchitect* MaxCAI, tAI, and GEMORNA mRNAs showed an intermediate persistence, while CUSTOM showed the lowest persistence.

We next tested whether sequence-based predictors of stability explained the observed intracellular persistence. We found that MFE was moderately correlated with intracellular stability (R²=0.392; **Figure 3c**), indicating that stable secondary structure moderately predicts intracellular stability. CAI and GC content were weakly correlated to intracellular stability (R²=0.233 and R²=0.228 respectively). This observation was sequence dependent, with significant differences in correlation observed between the two constructs (**Figure S7a, b**).

### Reactogenicity of *mRNArchitect* designs

The activation of the innate immune response by exogenous RNA is a key consideration in mRNA design and a challenge for long-term or repeated dosing. Cytoplasmic and endosomal RNA sensors recognise uncapped, double-stranded, or chemically unmodified RNA. Since *LinearDesign* introduces more double-stranded secondary structure, similar to DERNA (**Figure 4a; S8**), we next tested whether these design strategies differ in their activation of the innate immune response.

**Figure 4.**
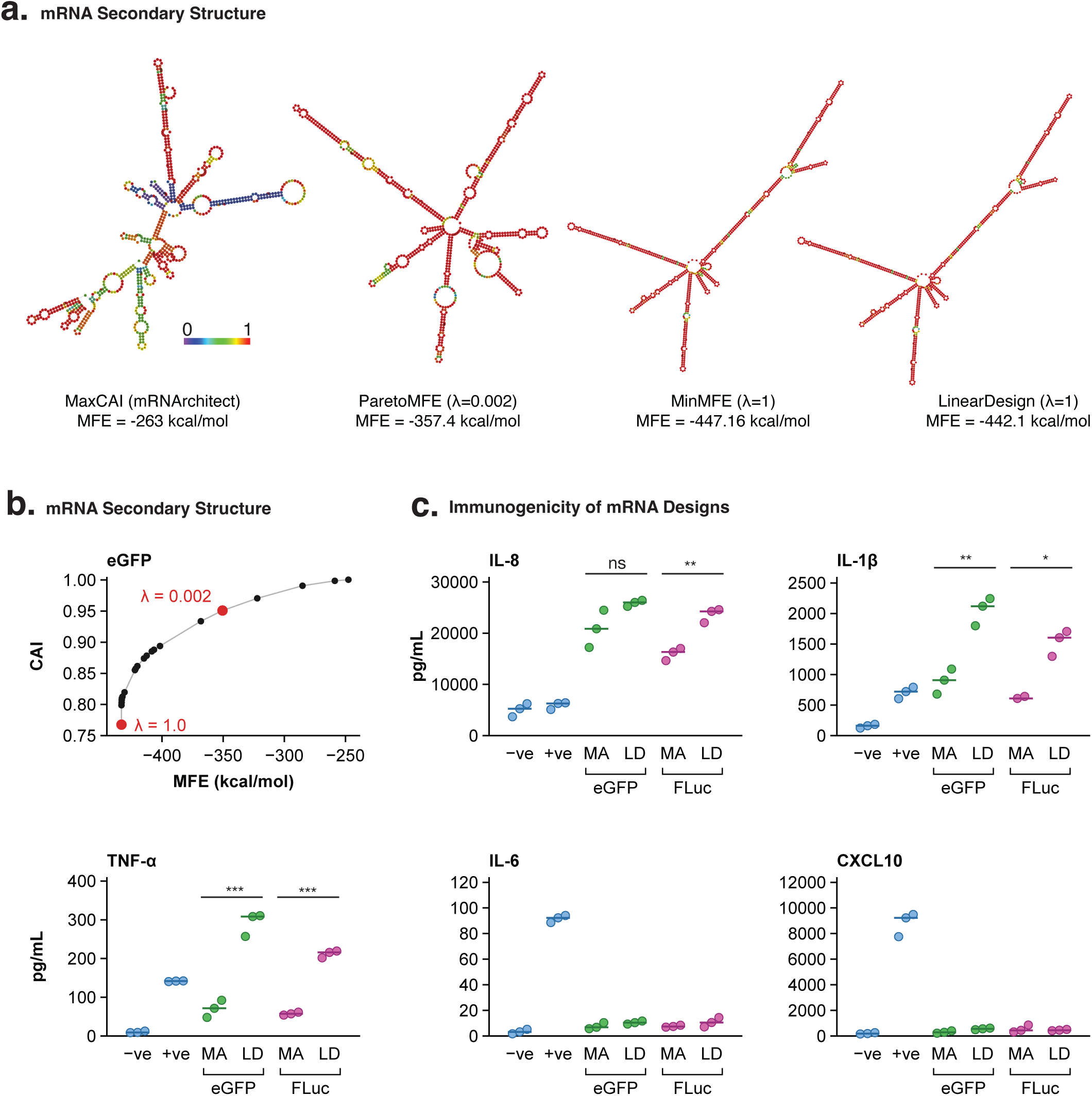
Predicted structure and innate immune activation of mRNA designs. (a) Predicted secondary structures of eGFP designs produced by MaxCAI, ParetoMFE (λ = 0.002), MinMFE (λ = 1) and LinearDesign, coloured by base-pairing probability, with minimum free energy below each. (b) CAI regressed against MFE across DERNA λ values for the eGFP coding sequence, with λ = 1.0 and λ = 0.002 marked. (c) Secretion of IL-8, IL-1β, TNF-α, IL-6 and CXCL10 following transfection of *mRNArchitect* (MA) and LinearDesign (LD) eGFP and luciferase mRNAs into differentiated macrophages, with untreated (−ve) and 3p-hpRNA (+ve) controls. Each point is one replicate; the bar is the median.

We assessed innate immune activation by measuring five cytokines (CXCL10, IL-6, IL-8, IL-1β, and TNF-α) following transfection of *LinearDesign* and *mRNArchitect MaxCAI FLuc* and *eGFP* mRNAs into a differentiated human monocyte-derived macrophage line (**Figure 4c**).

Across both eGFP and FLuc mRNA, *LinearDesign* constructs induced higher cytokine responses than their *mRNArchitect*-designed counterparts, with significantly higher IL-1β (Welch’s t-test; p=0.0029 and p<0.0159 respectively), and TNF-α (Welch’s t-test; p=0.0008 and p=0. 0003 respectively), consistent with NF-κB-driven transcriptional responses. Together, these results indicate that *mRNArchitect*-designed constructs elicit a lower inflammatory response than corresponding *LinearDesign* constructs.

### Untranslated regions selection and optimisation

The 5′ and 3′ UTRs are critical determinants of mRNA translation and stability, and *mRNArchitect* includes curated UTR libraries to support their selection (**Figure 1a**). By default, it uses 5′ and 3′ UTRs derived from the human and mouse α-globin genes (HBA1) respectively, which have demonstrated robust performance across a wide range of cell types and applications. To evaluate alternative UTRs, we screened a panel of 5′ UTR sequences using the FLuc reporter in HEK293T cells, including housekeeping gene UTRs (albumin, beta-actin), a minimal 5′ UTR, low-complexity UTRs (NELL2, ANKRD49, ZFAND6)^17^, and SCGB1A1, selected for its high, lung-restricted expression (**Table S4; Figure S9a**).

Relative to the default *mRNArchitect* HBA1 5′ UTR, beta-actin and SCGB1A1 modestly increased expression, while the minimal and albumin 5′ UTRs performed comparably (**Figure 5a**). Notably, the NELL2 5′ UTR, with a low GC content (56.1% in the UTR and cap-proximal CDS region; **Table S5**), drove the highest expression of all constructs tested. By contrast, ANKRD49 and ZFAND6 showed the lowest expression. Both secondary structure and GC content were positively correlated with FLuc expression at 24hpt (**Figure S10a–c**).

**Figure 5.**
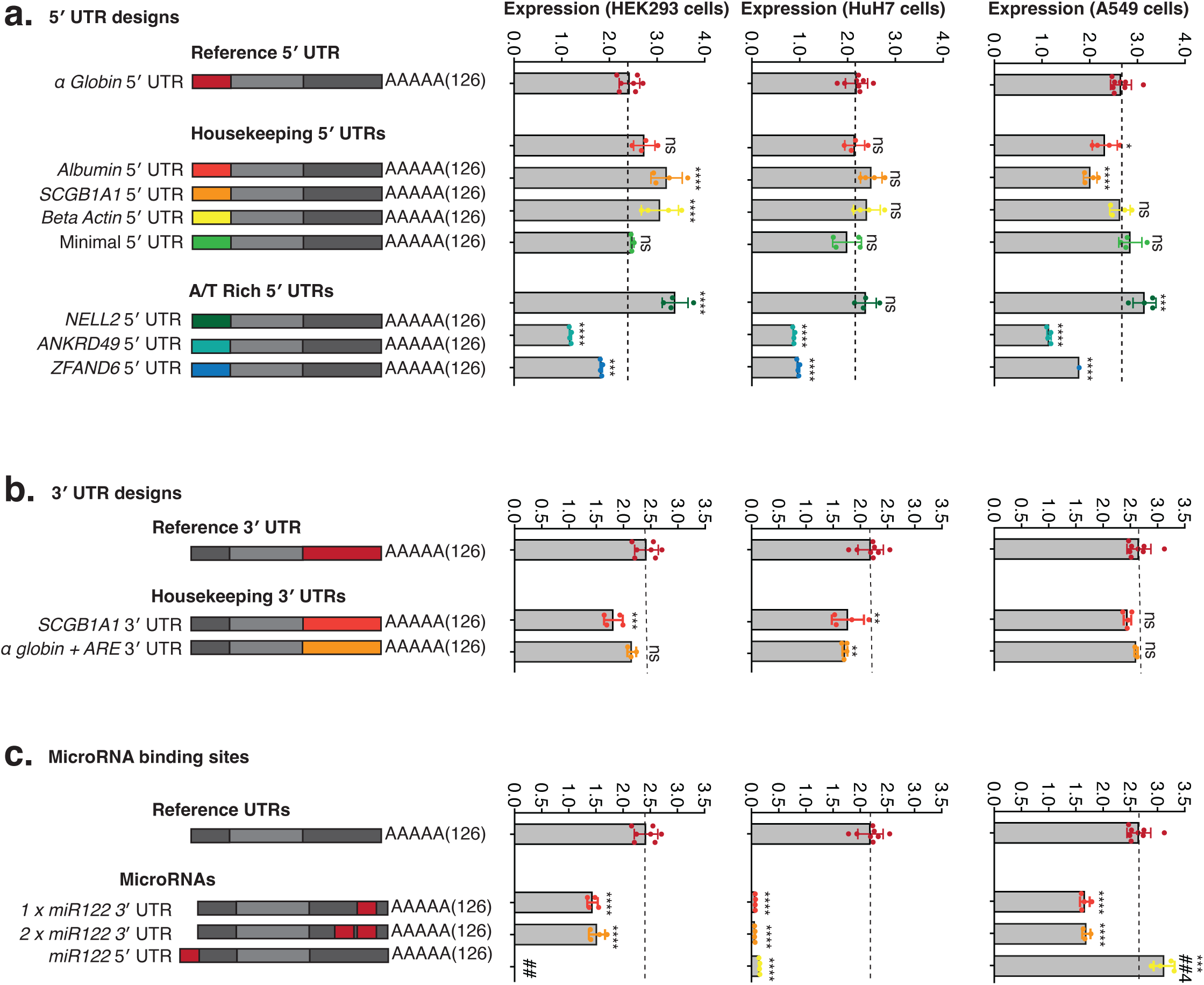
UTR and microRNA element designs. (a) Firefly luciferase expression from eight 5′ UTRs, comprising the α-globin reference, four housekeeping UTRs (albumin, SCGB1A1, beta-actin and a minimal UTR) and three A/T-rich UTRs (NELL2, ANKRD49, ZFAND6), in HEK293, Huh7 and A549 cells. (b) Expression from the α-globin reference 3′ UTR, the SCGB1A1 3′ UTR and an α-globin 3′ UTR carrying a 60 nt AU-rich element. (c) Effect of miR-122 binding sites inserted in the 3′ UTR, as one or two copies, or in the 5′ UTR, in the same three cell lines. Dashed line marks the reference construct.

We also evaluated two alternative 3′ UTR designs alongside the HBA1 α-globin default: a construct incorporating a 60-nucleotide AU-rich element (ARE60) positioned between the stop codon and the mouse α-globin 3′ UTR, previously shown to enhance mRNA stability and protein output^24^, and a construct incorporating the SCGB1A1 3′ UTR. Both designs showed modestly reduced FLuc expression relative to the α-globin control (**Figure 5b**).

The UTR panel showed broadly consistent results across further cell lines, including Huh7 (liver-derived) and A549 (lung-derived) (**Figure 5b**), highlighting the generalisability of these findings across cell lines. For example, the NELL2 5′ UTR consistently induced higher expression than the HBA1 5′ UTR, while ANKRD49 and ZFAND6 consistently induced lower expression. Moreover, the minimal 5′ UTR showed similar expression to HBA1 in all three cell lines.

Together, these findings establish a growing catalogue of experimentally validated UTRs for use in mRNA design. We anticipate this catalogue will continue to expand as additional regulatory elements are systematically screened and validated across further cell types and applications, providing an increasingly comprehensive, curated resource to inform UTR selection in *mRNArchitect*.

### microRNA binding sites

MicroRNAs bind complementary sequences within mRNAs and are key regulators of mRNA expression and stability within the cell. To avoid unwanted microRNA-mediated regulation, *mRNArchitect* can omit reverse complement sequences of selected microRNA seed sites that may otherwise reduce the performance of mRNA medicines.

Conversely, the incorporation of synthetic microRNA binding sites into mRNA medicines can provide tissue- or cell-type-specific control of expression. Accordingly, the UTR sequence input field includes a set of validated microRNA landing pads that enable synthetic microRNA binding sites to be incorporated into 5′ or 3′ UTRs (**Figure 1a**). We selected binding sites for miR-122, a liver-enriched microRNA that suppresses translation in liver cells with minimal impact on extrahepatic expression. One or two miR-122 binding sites were incorporated into either the 5′ or 3′ UTR of luciferase mRNAs (**Figure 5c**).

We tested the impact of these synthetic binding sites in Huh7 cells, which endogenously express high levels of miR-122, alongside human kidney-derived HEK293T and lung-derived A549 cells, which do not (**Figure 5c**). In Huh7 cells, all constructs containing miR-122 binding sites showed reduced expression relative to controls, confirming effective, functional suppression of liver-associated translation. In HEK293T and A549 cells, the 5′ UTR miR-122 construct showed little effect on expression, consistent with the absence of endogenous miR-122 in these cell types.

Together, these results establish a set of validated microRNA landing pads that can be used to incorporate defined microRNA binding sites into mRNA UTRs, enabling programmable, cell-type-specific control of expression.

## DISCUSSION

The optimisation of mRNA sequences can enhance translation and stability and reduce immunogenicity, and is a critical first step in the development of safe and effective mRNA medicines. Here we present *mRNArchitect*, software for the assembly and optimisation of mRNA medicines. By benchmarking *mRNArchitect* against alternative mRNA design tools across expression, stability and immunogenicity, we identify the design principles that underpin mRNA performance and the trade-offs between them.

The design of an mRNA sequence presents a choice between multiple strategies. Codon optimisation favours rapid and strong expression, whereas optimisation for secondary structure folds the mRNA into a compact structure that protects it from enzymatic degradation. However, the relative impact of each strategy on mRNA performance remains poorly understood.

We found that codon optimisation reliably generates the best-performing mRNAs. *mRNArchitect* codon-optimised designs showed the highest expression and the longest in-vial half-life of all designs tested. In contrast, structure-optimised designs produced lower expression and greater immunogenicity. Nonetheless, structured mRNAs persisted longer within the cell and showed a smaller relative decline in expression over time.

mRNA stability differed between the vial and the cell. This reflects distinct degradation processes, with physicochemical damage, such as hydrolysis and shearing in the vial, and enzymatic degradation by cellular nucleases within the cell. Notably, highly expressed mRNAs often showed the lowest intracellular persistence despite the highest in-vial stability, suggesting that rapid translation might increase the processing and turnover of mRNA within the cell. Longer intracellular persistence therefore did not compensate for lower expression. We recommend codon optimisation in most cases, while noting that secondary structure might offer an alternative design strategy for applications that require durable expression, and tolerate high reactogenicity.

AI models have recently emerged as an alternative strategy for mRNA design and are trained on large sequence datasets to learn the features associated with high expression. However, we found that the best-performing AI-based designs reached a similar sequence solution and experimental performance to codon optimisation, suggesting that these methods converge on a common solution.

Regardless of the design strategy, manufacturability is a critical consideration during sequence design. Dense GC regions and stable secondary structures can complicate DNA template synthesis and PCR amplification and can yield poor-quality mRNAs that undermine performance. Accordingly, structure-optimised sequences were more difficult to manufacture and yielded lower mRNA quality.

In addition to sequence optimisation, regulatory elements within an mRNA can control its expression and stability through complex interactions with cellular factors, including RNA-binding proteins, the translational machinery and microRNAs. Here we validated a range of UTRs and showed that engineered microRNA landing pads can program expression. As our understanding of cellular RNA biology improves and the elements that drive these processes are identified, they might be incorporated into mRNA designs to enable cell-type-specific control of stability, expression and subcellular localisation.

Together, these approaches can allow mRNA design to be tailored to the intended clinical application. Vaccines and gene editing might favour rapid, high-magnitude expression, whereas protein replacement therapies might favour slower, more durable expression. *mRNArchitect* supports this diversity through independent optimisation of sequence sub-regions, which allows users to balance these parameters and select the design strategy for each application. We also anticipate the provision of validated backbones and templates for key mRNA medicines.

In summary, codon optimisation reliably produced the best-performing mRNAs, and *mRNArchitect* provides a framework to apply this and other design strategies according to the intended application. These conclusions are based on two reporter mRNAs, and validation with additional therapeutic sequences will be required. We will continue to develop *mRNArchitect* based on ongoing experimental validations and our ever-evolving understanding of RNA biology, to support the development of new mRNA medicines.

## SUPPLEMENTARY FIGURES

**Supplementary Figure 1. mRNArchitect computational performance.** (a) Elapsed wall clock time and (b) peak memory usage for optimisation of 80 human coding sequences of 100 to 8,000 nt, using default parameters (blue) and constrained parameters (red). Each point is one sequence.

**Supplementary Figure 2. Sequence remodelling across mRNA classes.** Distribution of sequence properties for six mRNA classes before (wildtype, blue) and after (optimised, pink) *mRNArchitect* optimisation. Hamming ratio gives the proportion of nucleotide positions changed by optimisation, shown once per class. Remaining panels show Codon Stability Coefficient, predicted translation (RiboNN), predicted half-life, CAI, MFE, GC content and uridine depletion. Horizontal lines mark quartiles.

**Supplementary Figure 3. Benchmarking Luciferase mRNA sequences.** Sequence properties of Luciferase mRNAs optimised using different tools. Ten GEMORNA and CUSTOM designs were prepared using the same optimisation criteria, while ten LinearDesign sequences were prepared using different values of Lambda. A single design was prepared for all other tools.

**Supplementary 4. Quality control of manufactured mRNAs.** Capillary gel electrophoresis profiles of each mRNA used in this study, with the percentage of correctly sized products given for each. Nucleotide lengths indicated on the left.

**Supplementary Figure 5. Sequence features and in-vial mRNA stability.** Half-life under accelerated thermal degradation at 90°C plotted against CAI, GC content, MFE, predicted DegScore half-life, and tAI, for (a) luciferase and (b) eGFP mRNA designs. Dashed lines show linear regression with R².

**Supplementary Figure 6. Sequence features and intracellular mRNA stability.** Intracellular persistence, expressed as the ratio of transcript abundance at 6 h to 48 h post-transfection in HEK293T cells, plotted against CAI, GC content, MFE, predicted DegScore half-life, Codon Stability Coefficient and tAI, for (a) luciferase and (b) eGFP designs. Dashed lines show linear regression with R².

**Supplementary Figure 7. Sequence features and mRNA expression.** Expression at 24 h post-transfection in HEK293T cells plotted against CAI, MFE, GC content, tAI, Codon Stability Coefficient and predicted half-life, for (a) luciferase and (b) eGFP designs. Each point is one replicate. Dashed lines show linear regression with R².

**Supplementary Figure 8. Structural optimisation across representative sequences.** (a) Predicted secondary structures of firefly luciferase, eGFP, ovalbumin and SpCas9 coding sequences as wildtype and as optimised by MaxCAI, MinMFE and ParetoMFE (λ = 0.002), with MFE given below each. (b) CAI against MFE for the full range of DERNA λ values, with selected λ values labelled in red.

**Supplementary Figure 9. Expression profile of SCGB1A1 across human tissues.** Expression profiles of SCGB1A1 derived from Illumina RNA-seq, microarray, and SAGE datasets, obtained from the GeneCards database. All three datasets indicate strong and selective upregulation of SCGB1A1 in lung relative to all other tissues analysed.

**Supplementary Figure 10. 5′ UTR structure and composition determine expression.** (a) Predicted secondary structures spanning each 5′ UTR and the first 60 nt of the coding sequence, coloured by base-pairing probability. (b) Luciferase expression at 24 h against length-normalised MFE and (c) against GC content, in HEK293, Huh7 and A549 cells. Expression is normalised to the α-globin reference construct. Lines show linear regression with R².

## ACKNOWLEDGEMENTS

The authors acknowledge the facilities and the scientific and technical assistance of the BASE mRNA Facility at The University of Queensland. BASE is supported by Therapeutic Innovation Australia (TIA). TIA is supported by the Australian Government through the National Collaborative Research Infrastructure Strategy (NCRIS) program.

## SUPPLEMENTARY DATA

Supplementary data are available online.

## CONFLICT OF INTEREST

None.

## FUNDING

Data availability *mRNArchitect* is available through www.basefacility.org.au/software/ and can be downloaded for local installation from GitHub at https://github.com/BaseUQ/mRNArchitect

## METHODS

### *mRNArchitect* implementation and optimisation parameters

*mRNArchitect* is implemented in Python 3.13 and builds on the DNA Chisel constraint-solving framework (v3.2.16). It is available as a web application at https://basefacility.org.au/software/ and for local installation from https://github.com/BaseUQ/mRNArchitect. All analyses in this paper used version 1.4.1.

Users supply a coding sequence as amino acid, RNA or DNA. Lowercase input and uracil are normalised to uppercase DNA. A warning is given if the sequence does not begin with a methionine codon or end with at least one stop codon, and the sequence is rejected if it contains ambiguous residues or its length is not a multiple of three.

Codon optimisation is applied as a soft constraint, whereby the constraint score is maximised through iterative synonymous substitutions^14^. Depending on user input, codon selection is optimised according to either the Codon Adaptation Index (CAI)^5^, or the tRNA Adaptation Index (tAI)^15^.

Codon usage frequencies for CAI optimisation are obtained from the Kazusa Codon Usage Database (https://www.kazusa.or.jp/codon/ ), while tRNA gene copy numbers are retrieved from the Genomic tRNA Database (http://gtrnadb.ucsc.edu/ ). mRNArchitect calculates tAI using the optimised s-values, which represent the selective constraints on codon–anticodon coupling efficiency, as proposed by dos Reis et al (2003)^15^. CAI-based codon optimisation is supported for all organisms available in the Kazusa Codon Usage Database. In contrast, tAI-based optimisation is currently limited to human and mouse.

Manufacturability is enforced through hard constraints, which must be satisfied for a design to be returned. Global GC content is constrained by default to 0.40 to 0.70, and local GC content to 0.30 to 0.70 over a sliding 40 nt window, which resolves GC-rich regions that impede DNA synthesis. Homopolymer tracts are limited to eight consecutive A or U and five consecutive C or G residues, and all 10-mers must be unique. Hairpins are avoided using a minimum stem length of 20 nt evaluated over a 200 nt window. Users may exclude defined sequence strings, including restriction enzyme recognition sites, microRNA seed sites and ribosomal slip sequences^25^.

Secondary structure optimisation uses DERNA^16^, which applies dynamic programming to identify Pareto-optimal solutions balancing minimum free energy (MFE) against CAI. The parameter λ sets this balance, where λ = 1.0 weights the objective entirely towards MFE and λ = 0 entirely towards CAI. Secondary structure optimisation operates on the coding sequence alone and does not enforce the manufacturability constraints above. mRNArchitect incorporates different values of λ, which enable pareto-optimisation of mRNA sequences where CAI and secondary structure are differentially weighted.

Sub-region optimisation applies independent parameter sets to defined coordinate ranges, or excludes regions from optimisation to preserve validated linkers, signal peptides, siRNA binding sites and other functional elements. *mRNArchitect* then assembles the coding sequence with the selected UTRs and poly(A) tail and returns three candidate sequences with a design report recording the input, all parameters, the software version and the properties of each output.

To assess *mRNArchitect* computational performance, it was benchmarked using default and constrained codon optimisation parameters under a local installation. Default optimisation parameters were: organism = *Homo sapiens*, global GC content 0.4 – 0.7, 40nt windowed GC content 0.3 – 0.7, no uridine depletion, no specified avoid sequences, avoid repeat length >10, avoid poly(A/U) = 9; avoid poly(C/G) = 6, hairpin stem size 10, hairpin window 60. Constrained optimisation parameters were the same as default parameters except avoid repeat length >7 was used.

### mRNA sequence optimisation using alternative tools and strategies

To assess optimisation across representative mRNA classes, we assembled six sequence sets: antibodies, vaccine antigens, CAR-T constructs, cancer-associated genes, housekeeping genes and human open reading frames. Antibody sequences were obtained from IMGT^26^, cancer-associated genes from the COSMIC Cancer Gene Census^27^, housekeeping genes from Eisenberg and Levanon^28^, and human open reading frames from the CCDS database^29,30^. Antigen and CAR-T sequences were curated from clinical-stage constructs.

Firefly luciferase and eGFP coding sequences were optimised with *mRNArchitect* using four strategies: codon optimisation by CAI (MaxCAI) or tAI (MaxtAI), MFE minimisation by DERNA (MinMFE, λ = 1.0), and a Pareto-optimal balance of MFE and CAI (ParetoMFE, λ = 0.002). All designs used the human codon table and default manufacturability constraints.

The same protein sequences were optimised with alternative tools. LinearDesign (v1.0), which prioritises secondary structure, was run with λ spanning 0 to 1 and the human codon table. AI-based codon optimisation models include GEMORNA (v1.0), CodonTransformer (v1.2.0) and CUSTOM, which were run with default parameters for human expression. Web-hosted tools were accessed in March 2026. Ten different sequences were generated for CUSTOM and GEMORNA, and the sequences with the strongest predicted expression were selected for further analysis.

Each design was assembled with the human α-globin 5′ UTR, the mouse α-globin 3′ UTR and a 126 nt poly(A) tail for manufacture. All sequences are listed in Supplementary Table 3.

### Sequence analysis and predictive modelling

Nucleotide composition, GC content and uridine content were calculated directly from each coding sequence. The Codon Adaptation Index was calculated following Sharp and Li^5^, and the tRNA Adaptation Index using the weights of dos Reis et al.^15^, both with the human tables applied during optimisation, so that reported values match the design report.

Minimum free energy and secondary structure were predicted with ViennaRNA (v2.6.4)^31^ at 37°C using default parameters. Folding was performed on the coding sequence alone unless otherwise stated. MFE density was defined as MFE divided by sequence length, in kcal mol⁻¹ per nucleotide, to allow comparison between constructs of different length.

Sequence divergence was quantified as the Hamming distance ratio^32^, the proportion of nucleotide positions differing between two sequences of equal length, calculated between each optimised sequence and its wild-type counterpart and pairwise between designs.

Predicted translation efficiency was calculated with RiboNN (v1.0)^19^, and predicted stability with the Codon Stabilization Coefficient^18^, averaged across all codons, and DegScore (v1.0). DegScore half-life^8^ values are relative and are used only to rank designs.

Analyses were performed in Python 3.11 using Biopython, NumPy, SciPy and pandas, with code available at https://github.com/BaseUQ/mRNArchitect.

### DNA template preparation, in vitro transcription and mRNA quality control

Plasmid templates encoding each mRNA design were synthesised commercially, as described in Leighton et al. (2025)^33^. Each coding sequence was flanked by the human α-globin 5′ UTR and the mouse α-globin 3′ UTR. Templates were PCR amplified using a forward primer incorporating a modified T7 promoter (5′-TAATACGACTCACTATAAGG-3′) optimised for co-transcriptional CleanCap AG incorporation, and a reverse primer carrying a 126 nt poly(A) tail. Products were purified with the QIAquick PCR Purification Kit (Qiagen) and confirmed for size and purity by capillary gel electrophoresis and UV spectrophotometry respectively.

mRNA was in vitro transcribed with T7 RNA polymerase, following Henderson et al. (2021)^34^ and the manufacturer’s instructions (NEB, E2080S). Linear DNA template (50 µg/mL) was incubated at 37 °C for 3 h with 16 µg/mL T7 RNA polymerase (NEB, M0251), ribonucleotides (6 mM ATP, 5 mM CTP, 5 mM GTP; NEB, N0450), 5 mM N1-methylpseudouridine-5′-triphosphate (TriLink, TRN108110), transcription buffer (40 mM Tris·HCl pH 8.0, 16.5 mM magnesium acetate, 10 mM DTT, 20 mM spermidine, 0.002% v/v Triton X-100), 2 U/mL yeast inorganic pyrophosphatase (NEB, M2403) and 1000 U/mL murine RNase inhibitor (NEB, M0314). Cap-1 structures were incorporated co-transcriptionally by addition of 4 mM CleanCap AG (TriLink, TRN711310). Reactions were stopped with 200 U/mL DNase I (NEB, M0303) at 37°C for 15 min, and mRNA was purified with Monarch RNA Cleanup Kits (NEB, T2050) and eluted in 1 mM sodium citrate, pH 6.4.

mRNA yield and purity were assessed by UV spectrophotometry (NanoPhotometer N120, Implen). Integrity and size distribution were assessed by automated capillary gel electrophoresis on an Agilent TapeStation with RNA ScreenTape (Agilent, 5067-5576), with samples denatured at 72 °C for 3 min.

### Cell culture and transfection

HEK293T, Huh7 and A549 cells were obtained from ATCC and maintained in Dulbecco’s Modified Eagle Medium (Gibco) supplemented with 10% foetal bovine serum and 1% penicillin-streptomycin, at 37°C in 5% CO . A549 cells were maintained in F-12K medium with the same supplements. Cells were passaged at 80% confluency, used below passage 25, and confirmed free of mycoplasma.

Cells were seeded the day they were transfected and reverse transfected using Lipofectamine MessengerMAX (Invitrogen) according to the manufacturer’s instructions, with a Lipofectamine-only negative control included in every experiment. For luciferase assays, cells were seeded at 0.3 × 10^5^ cells/mL in 96-well plates and transfected with 0.3 µg firefly luciferase mRNA per 10 cells. For eGFP assays, cells were seeded at 0.27 × 10 cells/mL in 24-well plates and transfected with 1.85 µg eGFP mRNA per 10 cells. For RT-qPCR, cells were seeded and transfected in 24-well plates as described for the corresponding reporter. Cells were harvested at 6, 24 and 48 h post-transfection.

### Reporter expression assays

Firefly luciferase expression was measured at 6, 24 and 48 h post-transfection using the NanoGlo Dual Luciferase Reporter Assay System (Promega) according to the manufacturer’s protocol. Luminescence was read on an Infinite 200 Pro plate reader (Tecan). Four technical replicates were assayed per condition and timepoint.

eGFP expression was measured at the same timepoints by flow cytometry. Cells were harvested and resuspended in 200 µL ice-cold dPBS containing 0.03 µg/mL Fixable Viability Stain 700 (BD Horizon), incubated on ice in the dark for 30 min, and analysed on a CytoFLEX flow cytometer (Beckman Coulter). At least 10,000 events were recorded per sample. Cells were gated sequentially on forward and side scatter, singlets, and viability stain-negative cells, and mean fluorescence intensity was calculated for the viable population. Four technical replicates were assayed per condition and timepoint.

To screen UTR variants, firefly luciferase constructs carrying each candidate 5′ or 3′ UTR were transfected into HEK293T, Huh7 and A549 cells and assayed at 24 h as described above for Firefly Luciferase. Constructs carrying one or two miR-122 binding sites in the 3′ UTR, or one site in the 5′ UTR, were assayed in parallel in the same three cell lines.

Expression is reported relative to the α-globin UTR reference construct assayed on the same plate.

### mRNA stability analysis

In-vial stability was assessed under accelerated thermal stress. Aliquots of 2 µg of each mRNA were incubated at 90 °C for 0, 20, 30, 40, 50, 60, 70, 80 and 90 min, then transferred to ice. Integrity was measured by automated capillary gel electrophoresis on an Agilent Fragment Analyser using the HS mRNA kit and recorded as the percentage area under the curve within the expected size range. Values were normalised to the 0 min control for each mRNA. Degradation was modelled as one-phase exponential decay, Y = 100e^(−KX), where Y is percentage integrity at time X and K is the decay rate constant, with the initial value constrained to 100% and the plateau to zero. Half-life was calculated as ln(2)/K.

Intracellular stability was assessed by RT-qPCR. HEK293T cells were transfected with firefly luciferase or eGFP mRNA as described above and harvested at 6, 24 and 48 h, with three technical replicates per condition. Total RNA was extracted with the Monarch Spin RNA Isolation Kit (NEB, T2110), and 10 ng was reverse transcribed with LunaScript RT SuperMix (NEB, M3010E). cDNA was diluted 1:10 and amplified with Luna Universal Probe qPCR Master Mix on a CFX Opus 96 system (Bio-Rad), using 95°C for 60 s followed by 45 cycles of 95°C for 15 s and 60°C for 30 s. Both reporters were detected with a single FAM-labelled probe and primer pair targeting the mouse α-globin 3′ UTR common to all constructs, multiplexed with a HEX-labelled GAPDH probe as the reference gene (**Table S6**). Amplification efficiencies were 90% and 93% for the luciferase constructs and 90% and 92% for the eGFP constructs. Relative transcript abundance was calculated by the comparative 2^(−ΔΔCq) method^35^. Intracellular persistence was expressed as the ratio of abundance at 6 h to that at 48 h.

### Innate immune activation

Innate immune activation was assessed by a commercial provider (RNAte) using their established protocol. Briefly, THP-1 monocytes were differentiated into macrophages with 100 ng/mL phorbol 12-myristate 13-acetate for 48 h, rested for 24 h, and transfected with 0.5 µg of each candidate mRNA. Transfection with 3p-hpRNA served as a positive control and untreated cells as a negative control. Supernatants were collected at 24 h and assayed for CXCL10, TNF-α, IL-6, IL-8 and IL-1β by ELISA and cytometric bead array. Assays were performed in triplicate across two independent experiments.

### Statistical analysis

All statistical analyses were performed in GraphPad Prism v11.0.0. Data are presented as mean ± standard deviation of four technical replicates for reporter expression assays and three technical replicates for RT-qPCR, unless otherwise stated. Comparisons between design tools were made by one-way ANOVA with Dunnett’s correction for multiple comparisons against the *mRNArchitect* MaxCAI design. UTR and microRNA constructs were compared against the α-globin UTR reference construct by the same method. Cytokine responses were compared by Welch’s t-test. Significance is indicated as ns (not significant), *P < 0.05, **P < 0.01, ***P < 0.001 and ****P < 0.0001. Relationships between sequence properties and measured expression or stability were assessed by ordinary least squares linear regression, with the coefficient of determination (R²) reported.

## Notes

### Competing Interest Statement

The authors have declared no competing interest.

### Summary of Updates

The version of this manuscript has been revised to include the description of new optimisaiton algorithms incorporated in our software, and further benchmarking results.

