## Supplementary Figures for "mRNArchitect: sequence design of mRNA medicines"

#### FIGURE S1

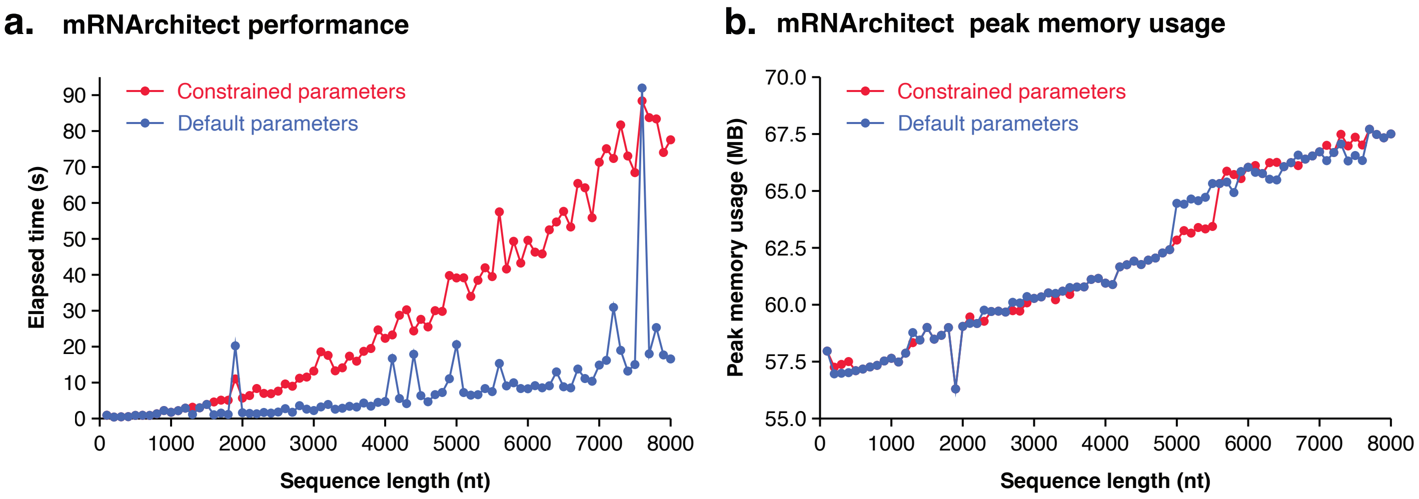
a.
mRNArchitect performance
Constrained parameters
Default parameters
Elapsed time (s)
Sequence length (nt)
b.
mRNArchitect peak memory usage
Constrained parameters
Default parameters
Peak memory usage (MB)
Sequence length (nt)

### FIGURE S2

**a.** mRNA sequences

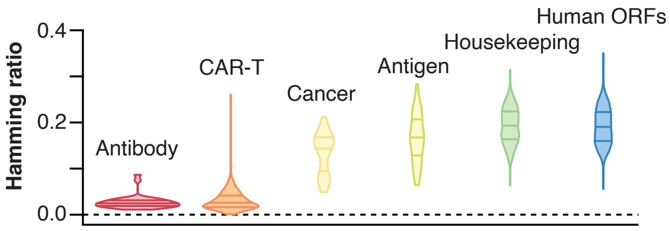


**b.** mRNA sequence remodeling

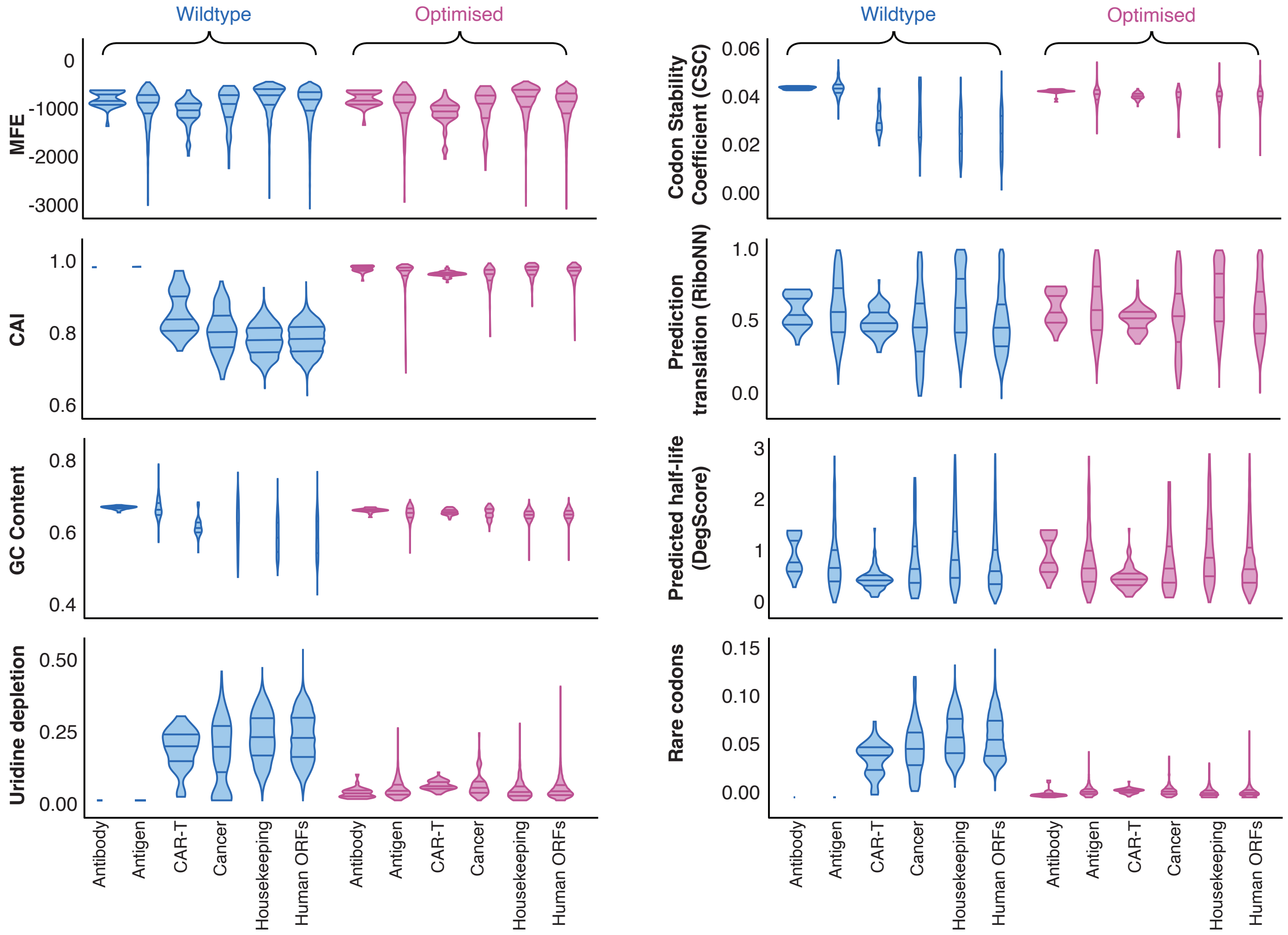

### FIGURE S3

**a.** Sequence features

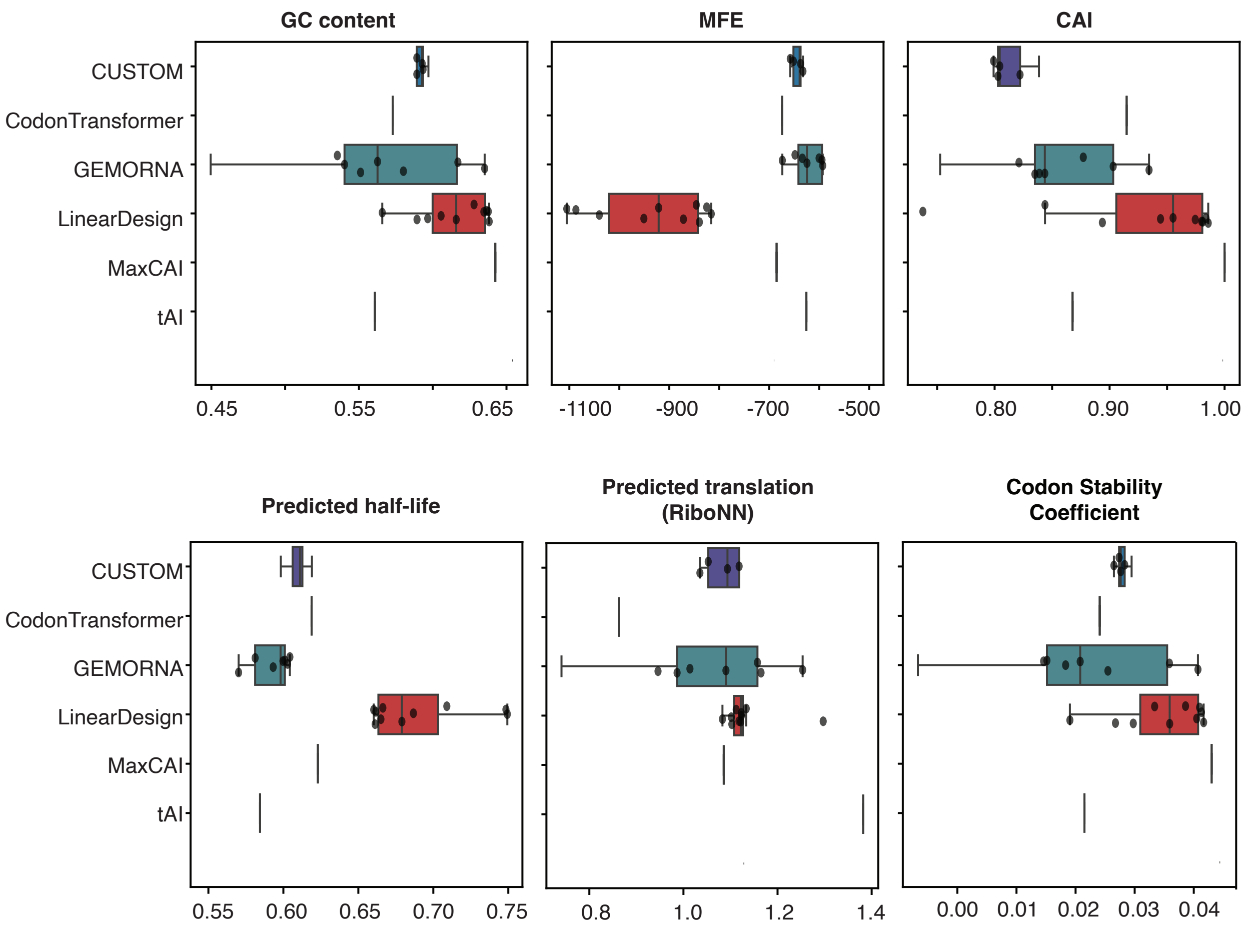

### FIGURE S4

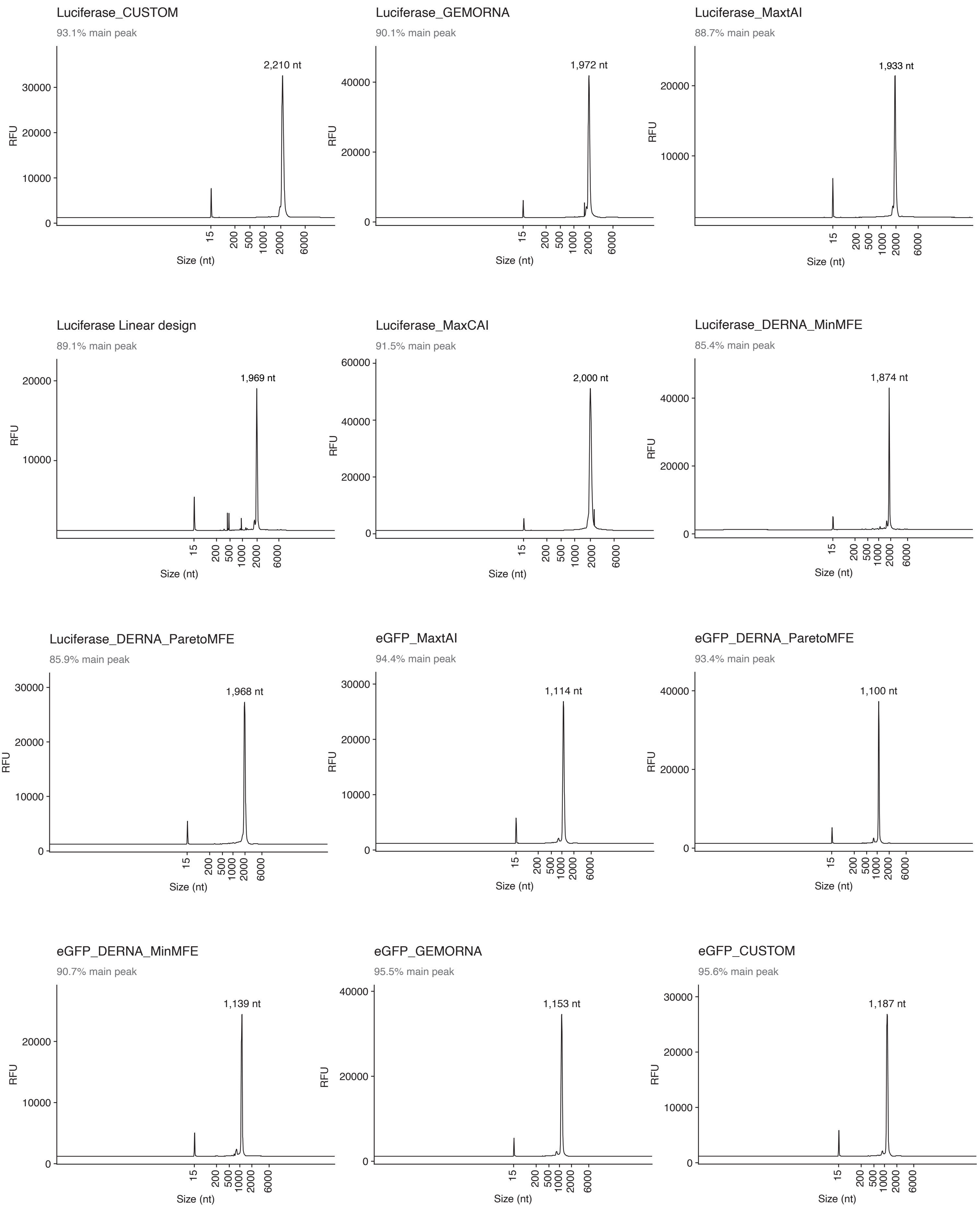
Luciferase_CUSTOM
93.1% main peak
2,210 nt
Luciferase_GEMORNA
90.1% main peak
1,972 nt
Luciferase_MaxtAI
88.7% main peak
1,933 nt
Luciferase Linear design
89.1% main peak
1,969 nt
Luciferase_MaxCAI
91.5% main peak
2,000 nt
Luciferase_DERNA_MinMFE
85.4% main peak
1,874 nt
Luciferase_DERNA_ParetoMFE
85.9% main peak
1,968 nt
eGFP_MaxtAI
94.4% main peak
1,114 nt
eGFP_DERNA_ParetoMFE
93.4% main peak
1,100 nt
eGFP_DERNA_MinMFE
90.7% main peak
1,139 nt
eGFP_GEMORNA
95.5% main peak
1,153 nt
eGFP_CUSTOM
95.6% main peak
1,187 nt
RFU
Size (nt)
15
200
500
1000
2000
6000

#### FIGURE S5

**a.** **Luciferase mRNA expression at 24 h**

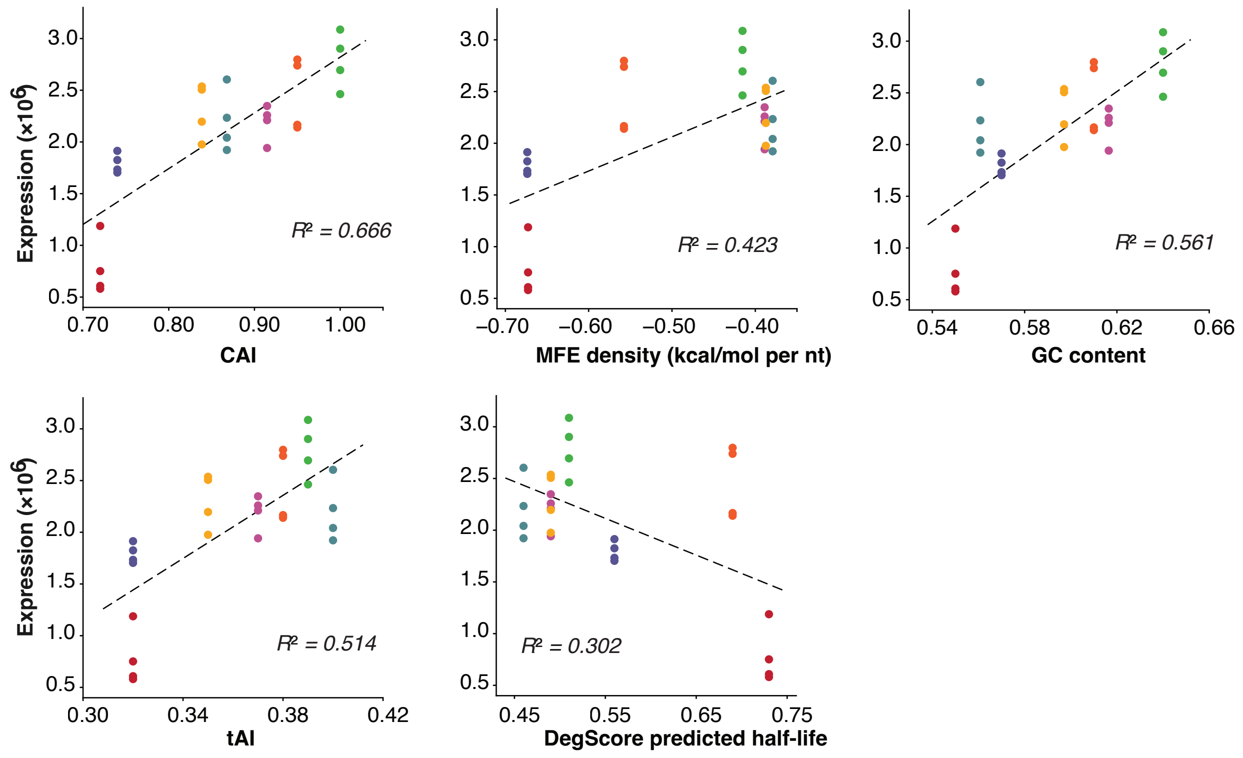


**b.** **eGFP mRNA expression at 24 h**

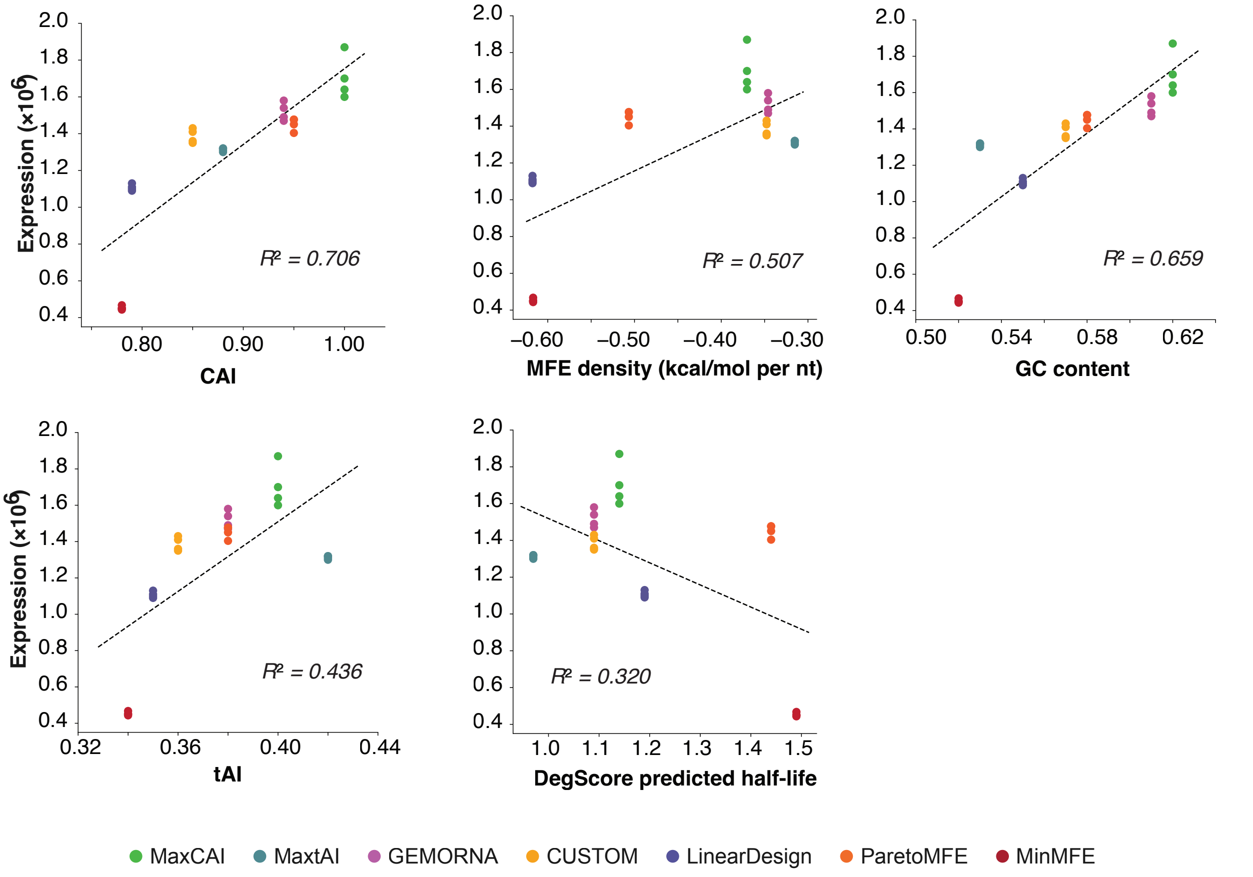

FIGURE S6

**a.** Luciferase mRNA in vial stability

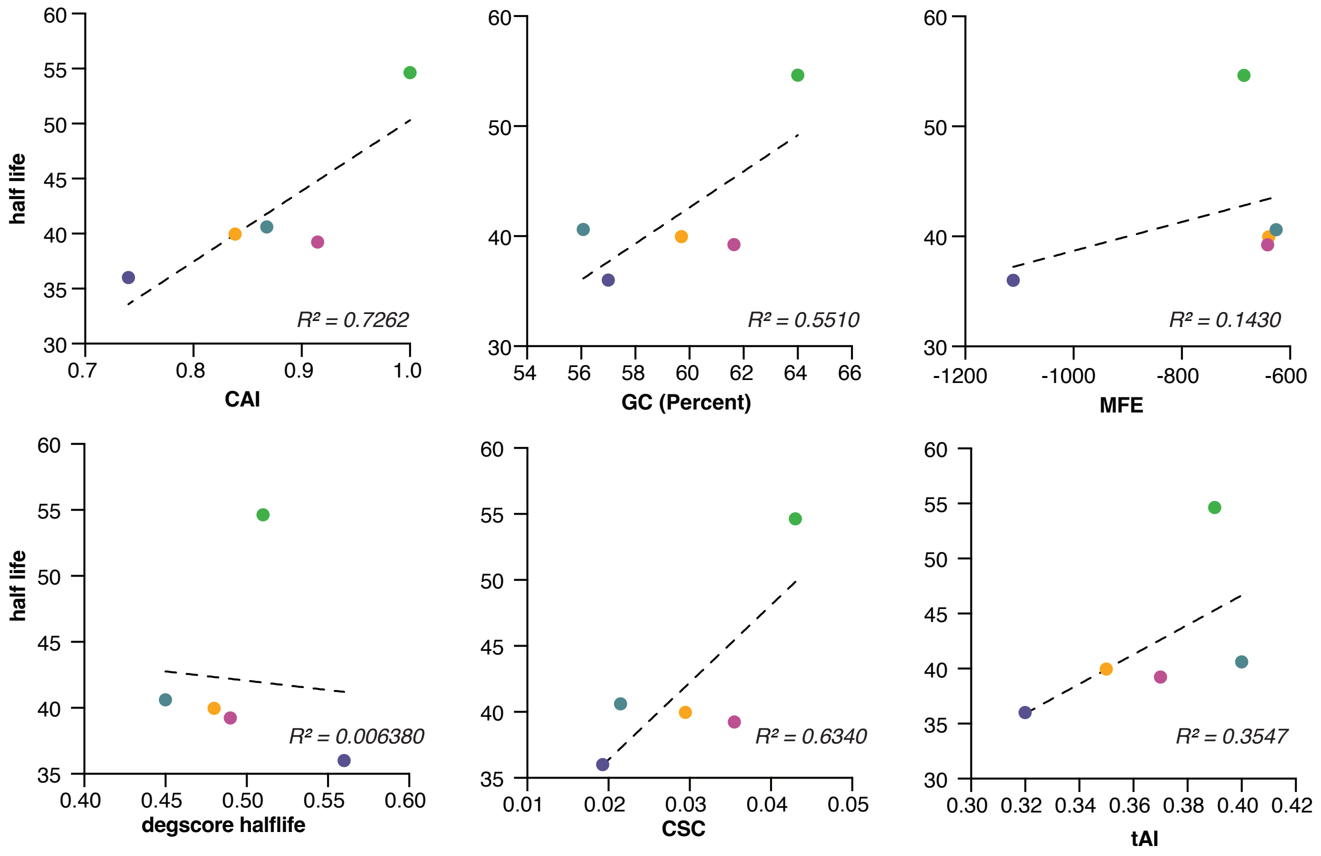


**b.** eGFP mRNA in vial stability

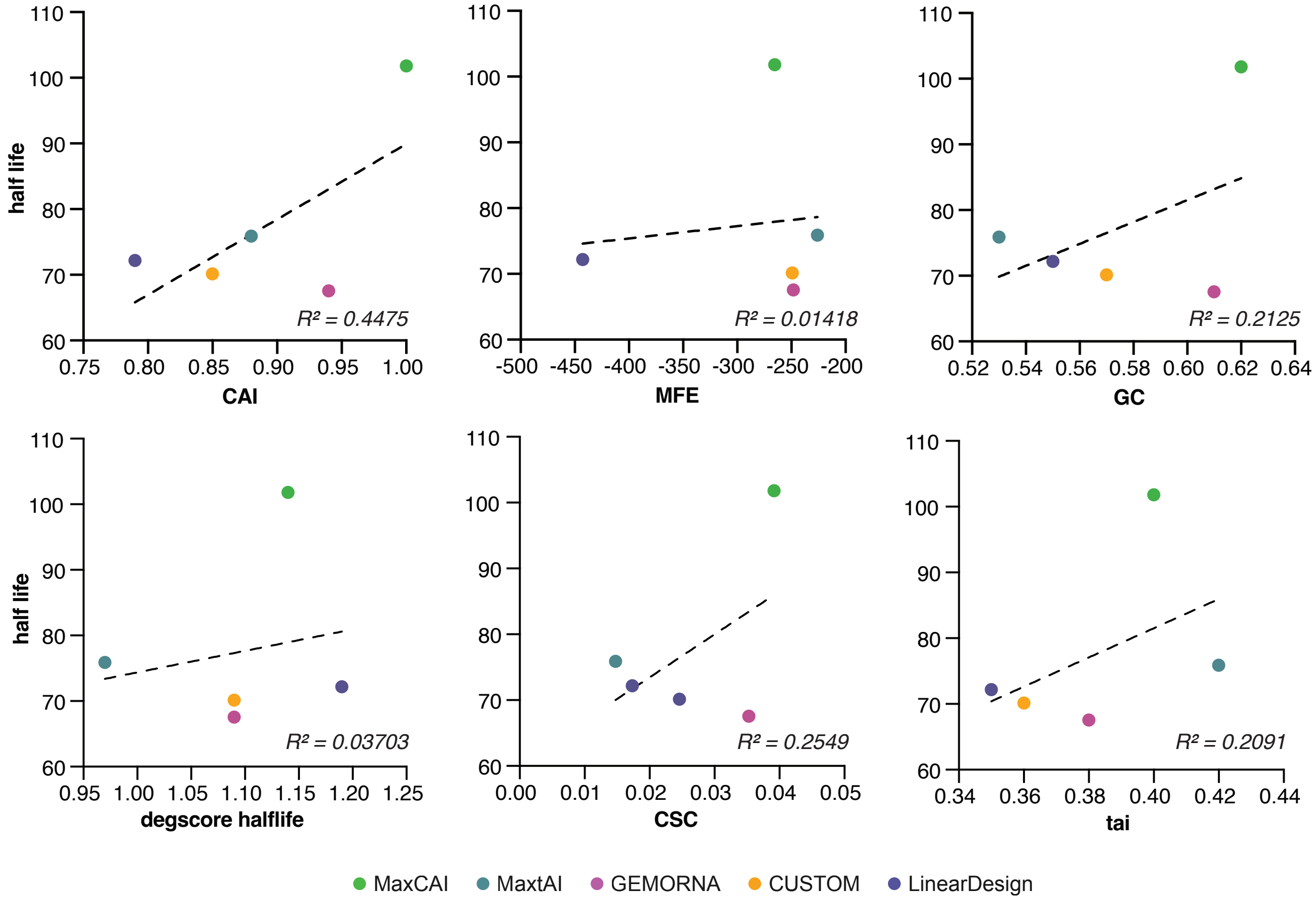

### FIGURE S7

**a.** Luciferase mRNA intracellular stability

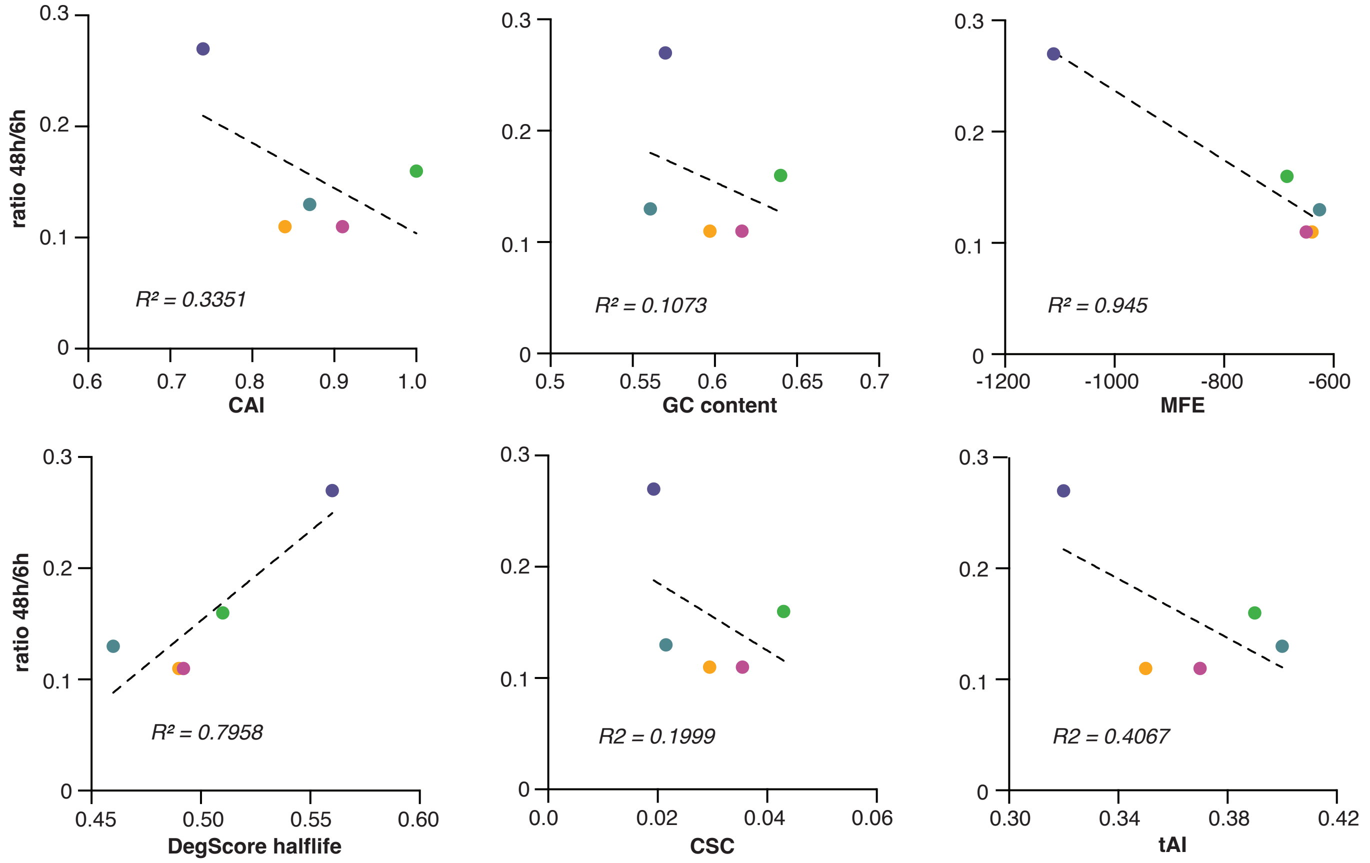


**b.** eGFP mRNA intracellular stability

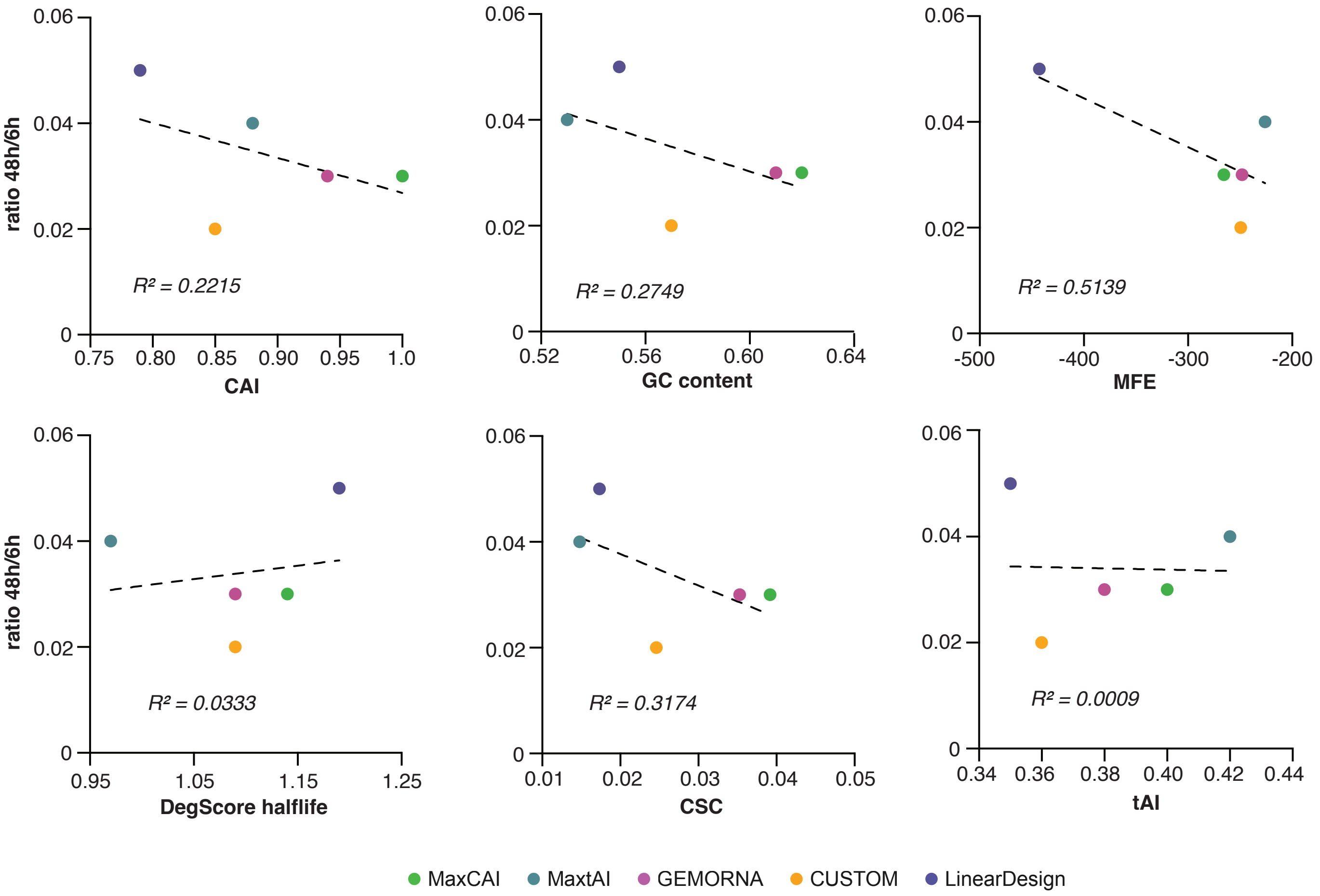

### FIGURE S8

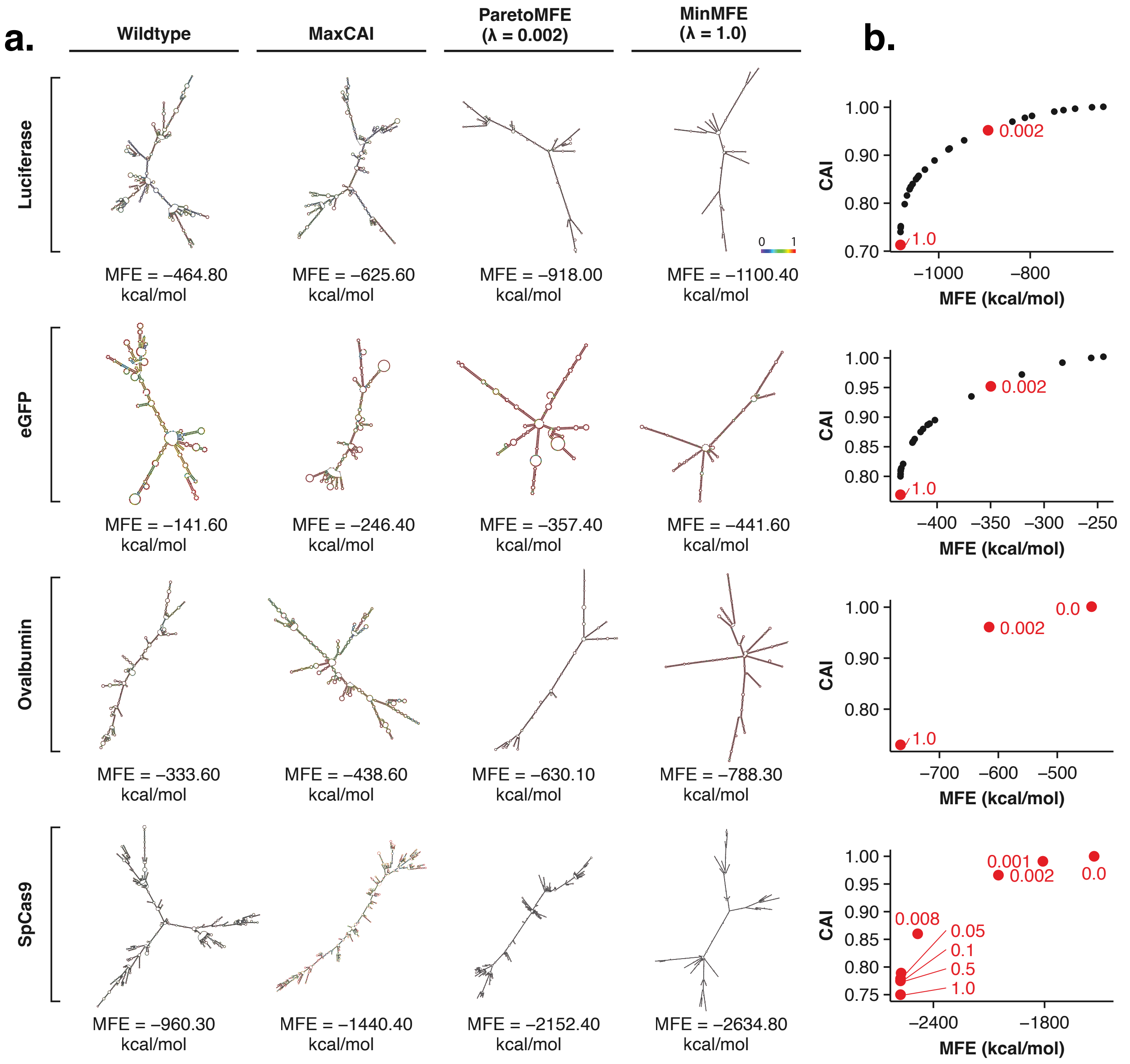
a.
Wildtype
MaxCAI
ParetoMFE (λ = 0.002)
MinMFE (λ = 1.0)
Luciferase
MFE = −464.80 kcal/mol
MFE = −625.60 kcal/mol
MFE = −918.00 kcal/mol
MFE = −1100.40 kcal/mol
0
1
eGFP
MFE = −141.60 kcal/mol
MFE = −246.40 kcal/mol
MFE = −357.40 kcal/mol
MFE = −441.60 kcal/mol
Ovalbumin
MFE = −333.60 kcal/mol
MFE = −438.60 kcal/mol
MFE = −630.10 kcal/mol
MFE = −788.30 kcal/mol
SpCas9
MFE = −960.30 kcal/mol
MFE = −1440.40 kcal/mol
MFE = −2152.40 kcal/mol
MFE = −2634.80 kcal/mol
b.
CAI
MFE (kcal/mol)
0.002
1.0
0.0
0.001
0.008
0.05
0.1
0.5

**a.** **mRNA expression in Normal Tissues for SCGB1A1 Gene**

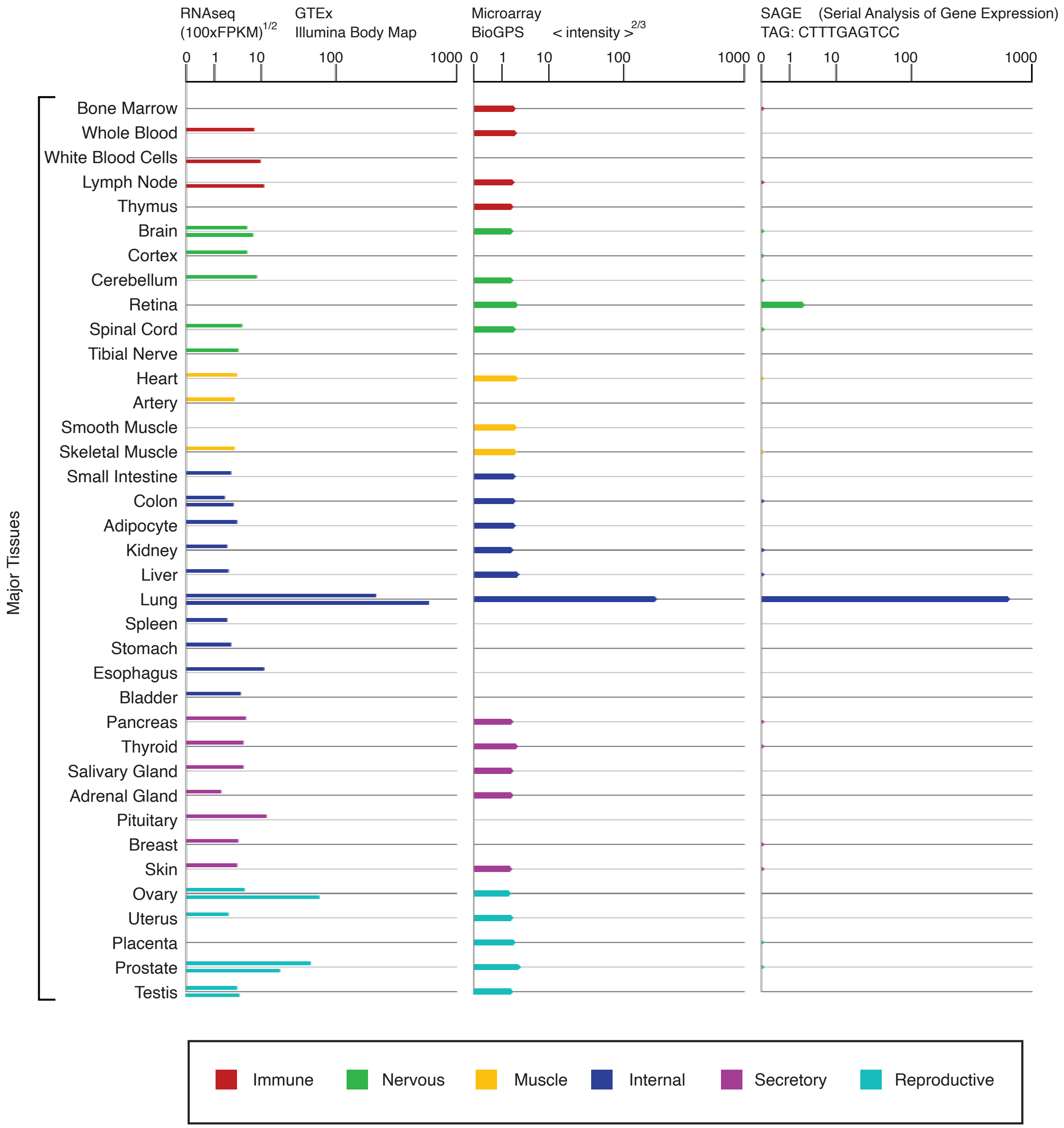

FIGURE S10

**a.** **Secondary structures proximal to the start codon**

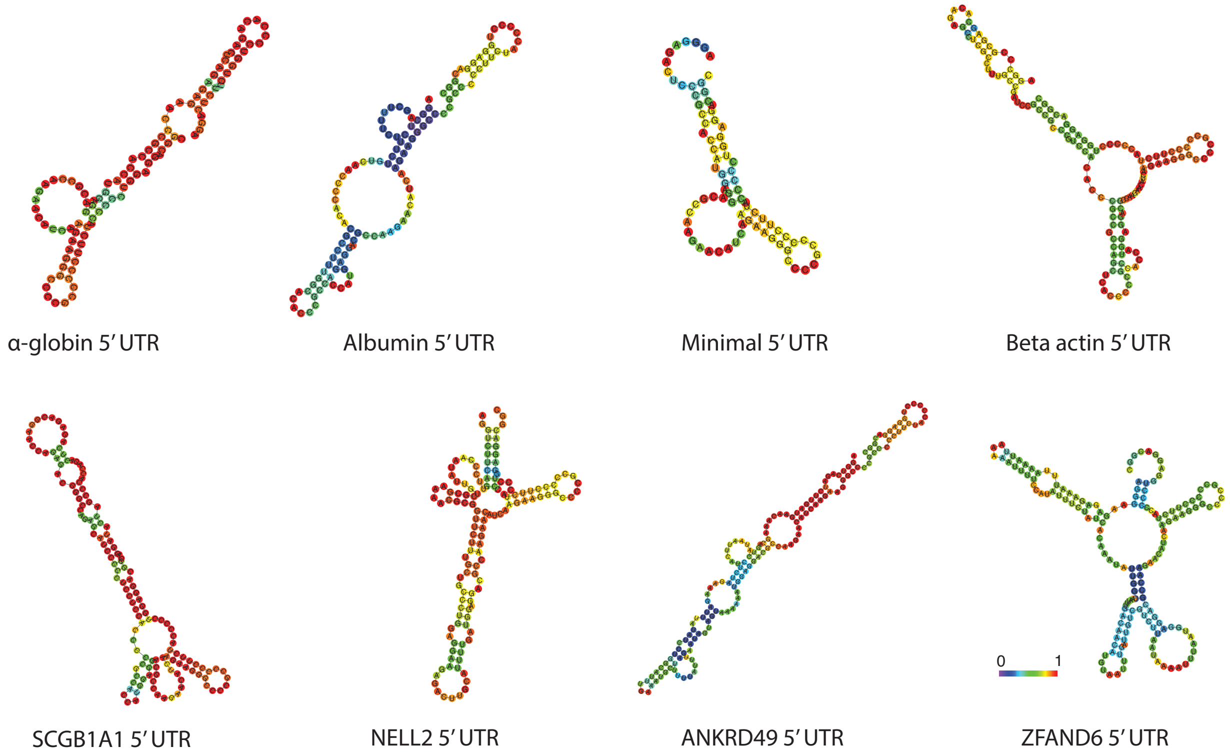


**b.** **Impact of 5′ UTR secondary structure on expression**

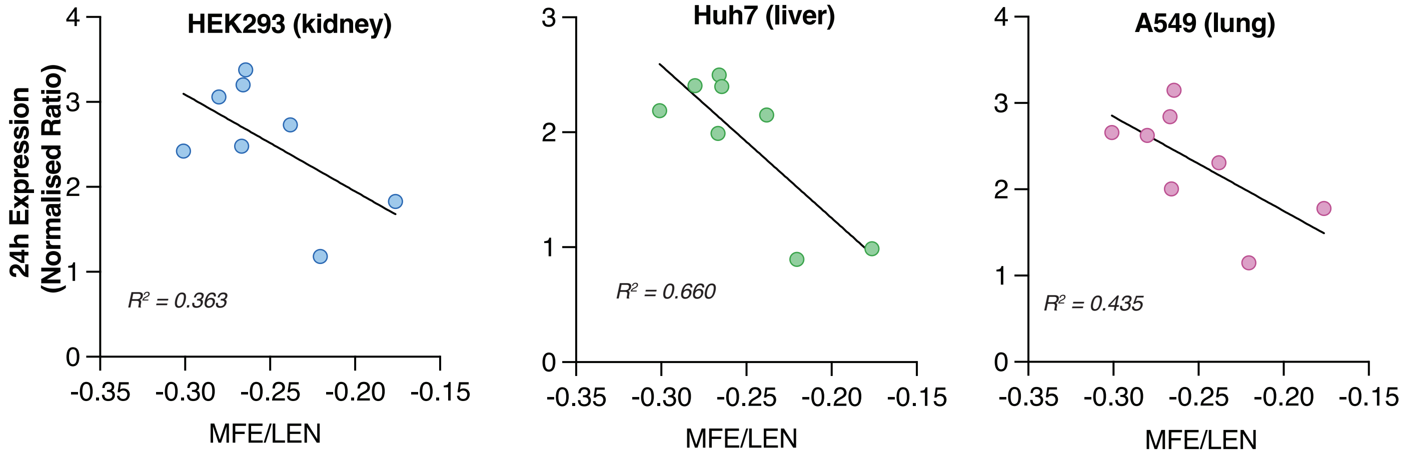


**c.** **Impact of 5′ UTR GC content on expression**

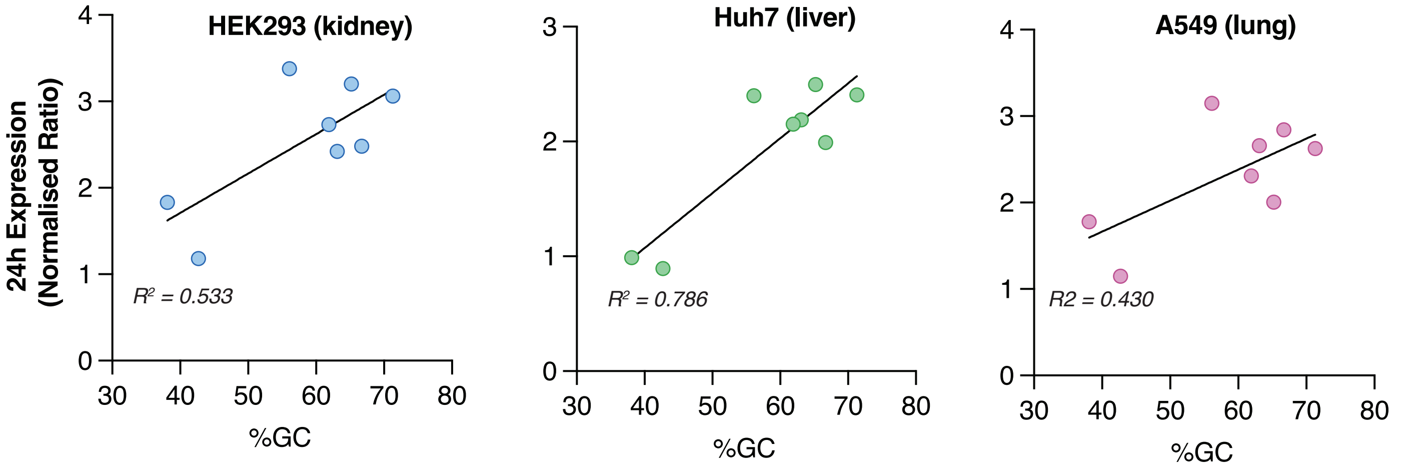
